# Shiga toxin remodels the intestinal epithelial transcriptional response to Enterohemorrhagic Escherichia coli

**DOI:** 10.1101/2020.08.11.245555

**Authors:** Alyson R. Warr, Carole J. Kuehl, Matthew K. Waldor

## Abstract

Enterohemorrhagic *Escherichia coli* (EHEC) is a food-borne pathogen that causes diarrheal disease and the potentially lethal hemolytic uremic syndrome. We used an infant rabbit model of EHEC infection that recapitulates many aspects of human intestinal disease to comprehensively assess colonic transcriptional responses to this pathogen. Cellular compartment-specific RNA-sequencing of intestinal tissue from animals infected with EHEC strains containing or lacking Shiga toxins (Stx) revealed that EHEC infection elicits a robust response that is dramatically shaped by Stx, particularly in epithelial cells. Many of the differences in the transcriptional responses elicited by these strains were in genes involved in immune signaling pathways, such as *IL23A,* and coagulation, including *F3*, the gene encoding Tissue Factor. RNA FISH confirmed that these elevated transcripts were found almost exclusively in epithelial cells. Collectively, these findings suggest that within the intestine, Stx primarily targets epithelial cells, and that the potent Stx-mediated modulation of innate immune signaling skews the host response to EHEC towards type 3 immunity.

**Significance Statement:** Enterohemorrhagic *Escherichia coli* (EHEC) is a potentially lethal foodborne pathogen. During infection, EHEC releases a potent toxin, Shiga toxin (Stx), into the intestine, but there is limited knowledge of how this toxin shapes the host response to infection. We used an infant rabbit model of infection that closely mimics human disease to profile intestinal transcriptomic responses to EHEC infection. Comparisons of the transcriptional responses to infection by strains containing or lacking Stx revealed that this toxin markedly remodels how the epithelial cell compartment responds to infection. Our findings suggest that Stx biases the intestinal innate immune response to EHEC and provide insight into the complex host-pathogen dialogue that underlies disease.

## Introduction

Enterohemorrhagic *Escherichia coli* (EHEC) is an important foodborne pathogen responsible for up to 2 million annual cases of acute gastrointestinal illness (1, 2). Chiefly colonizing the colon, EHEC typically leads to self-limiting hemorrhagic colitis; however, 5-10% of infected individuals also develop hemolytic uremic syndrome (HUS), a potentially life-threatening complication that can lead to renal failure (3, 4). Supportive rehydration therapy remains the primary treatment for EHEC infection, while antibiotics are associated with elevated frequencies of HUS and therefore contraindicated (5). There is no vaccine available (6). EHEC infection is associated with an inflammatory response in the colon, with patients demonstrating elevated fecal leukocytes and calprotectin levels, indicative of neutrophilic colitis. Colonic biopsy samples from patients with EHEC infection exhibit inflammation, including edema, fibrin deposition, and neutrophil invasion, as well as hemorrhage (7–11).

*E. coli* O157:H7 is the most common EHEC serotype, but several other EHEC serotypes have been described. All serotypes classified as EHEC share two primary virulence factors: the ‘LEE’ pathogenicity island that encodes a type III secretion system (T3SS), and prophages that encode one or more Shiga toxins (12–14). The activities of EHEC’s T3SS effector proteins mediate the pathogen’s tight adherence to the colonic mucosa (15), and can promote or antagonize the inflammatory response in epithelial cells (16–20).

During infection, EHEC produces and releases one or more Shiga toxins (Stx) into the intestinal lumen. Stxs are potent AB_5_ subunit exotoxins which are thought to bind to the host cell surface glycosphingolipid globotriaosylceramide (Gb3). Once internalized into the eukaryotic cell cytosol, Stx catalyzes a site-specific depurination of the 28s rRNA, which leads to inhibition of protein synthesis and triggers the ribotoxic stress response, production of cytokines, and cell death (21–27). Absorption of Stx into the blood results in its systemic circulation and damage to endothelial cells, particularly in the renal microvasculature, leading to the characteristic findings of HUS, including renal failure and hemolysis.

The role of Stx in EHEC pathogenicity in the colon is controversial. Although Stx has been associated with colonic pathology (7, 8, 28–30), the mechanisms that explain these observations are unclear because the presence of Gb3 in the colonic epithelium has been disputed (31–33, 29). Some suggest that Stx does not directly act on the colonic epithelium, but instead transverses through the epithelial layer to primarily act on endothelial and immune cells (34–38). Others have suggested that Stx can enter colon epithelial cells by binding to alternative receptors, such as Gb4 (33, 39). Despite the ambiguity surrounding the mechanisms by which Stx exerts toxic effects in the colon during EHEC infection, purified Stx can stimulate inflammatory responses in cultured cells (40–43, 22, 29, 44–47, 26, 25). Analyses of the transcriptional responses to EHEC infection in cultured epithelial cells and organoids have also demonstrated that processes linked to inflammatory signaling, cytoskeletal organization, apoptosis, and stress responses are altered (48–51). However, to date, our knowledge of the host response to EHEC infection is almost exclusively derived from tissue-culture based studies. Because mice do not develop overt diarrhea or colonic pathology during EHEC infection (52, 53), no comprehensive in vivo analyses of how EHEC modifies colonic gene expression patterns during infection have been reported. Moreover, the extent to which Stx contributes to such gene expression changes in vivo is unclear.

Immune responses to bacterial pathogens have been grouped into type 1 or type 3 immunity (54). Type 1 responses are usually triggered by intracellular pathogens, chiefly through IL-12 stimulated IFNγ-producing cells, which lead to activation of cytotoxic lymphocytes that can destroy infected cells. Type 3 responses are usually triggered by extracellular pathogens. Typically, exposure to pathogen-associated molecular patterns (PAMPs) stimulates dendritic cells to produce IL1 and IL23, which stimulate immune effector cells to produce IL17 and IL22. These cytokines lead to changes in epithelial cells, promoting antimicrobial peptide production and barrier maintenance, and trigger infiltration of phagocytic cells to engulf extracellular pathogens. However, the associations of type 1 and type 3 immunity with intracellular and extracellular pathogens respectively are not strict. For example, *Citrobacter rodentium,* an extracellular pathogen that encodes a T3SS highly similar to EHEC’s, elicits a type 1 immune response (55–57); it is unknown if the same is true for Stx-producing EHEC.

Here, we used infant rabbits, a small animal model of EHEC infection where orogastric inoculation of the pathogen leads to a disease that closely mimics the intestinal manifestations of human EHEC disease (28, 30, 58), to investigate how Stx production in the gut modifies the cellular response of the colonic mucosa during EHEC infection. We compared the colonic epithelial and lamina propria transcriptional responses to WT and mutant EHEC lacking Stx genes. Collectively, our findings provide a comprehensive profile of the colonic transcriptional responses to EHEC and suggest that Stx, which primarily impacts the gene expression of epithelial cells, shifts innate immune responses to the pathogen from type 1 toward type 3 immunity.

## Results

### Shiga toxin promotes apoptosis and hemorrhage in the colonic mucosa

We used EDL933, a prototypical *E. coli* O157:H7 clinical isolate (referred to as WT below) (59, 60), to investigate how Shiga toxins modify the response of the colonic mucosa to EHEC infection. This strain encodes two Stx variants, Stx1 and Stx2. Loci encoding both variants were deleted to yield a strain referred to as ΔΔ*stx*. WT and ΔΔ*stx* were orogastrically administered to infant rabbits to determine if Stx influences EHEC intestinal colonization or the colonic mucosal response to infection. As described previously in experiments with a different EHEC clinical isolate (28), the burden of WT and ΔΔ*stx* in the colon did not differ (Fig S1), suggesting that Stx does not alter EHEC colonization in this model. Nonetheless, Stx appeared to contribute to the development of diarrhea as described (28).

Histopathologic analyses of colon samples from animals inoculated with WT, ΔΔ*stx*, or PBS (mock) were carried out at peak colonization (36-40 hours post inoculation). Compared to mock-treated control animals, colon samples from both WT and ΔΔ*stx* infected rabbits had prominent pathologic changes in the mid and distal colon, sites of maximal colonization. WT and ΔΔ*stx* infections led to similar levels of overall colonic inflammation, characterized by an increased number of small mononuclear cells in the submucosal tissue, as well as comparable levels of heterophil (lapine neutrophil) infiltration into the lamina propria and epithelium (Fig S2). Heterophils were occasionally observed in large rafts in the colonic lumen in samples from animals infected with either WT or ΔΔ*stx,* and both WT and ΔΔ*stx* infection elicited epithelial sloughing in the mid-colon (Fig S3). However, compared to the colonic pathology associated with the ΔΔ*stx* strain, there was significantly more apoptosis, indicated by widespread fragmented nuclei, and edema/hemorrhage, indicated by widespread blood and fluid accumulation in tissue, observed in samples from animals infected with the WT strain (Fig 1). These observations support the hypothesis that EHEC production of Stx in the intestine provokes local pathology including apoptosis and hemorrhage in the colonic mucosa.

**Figure 1:**
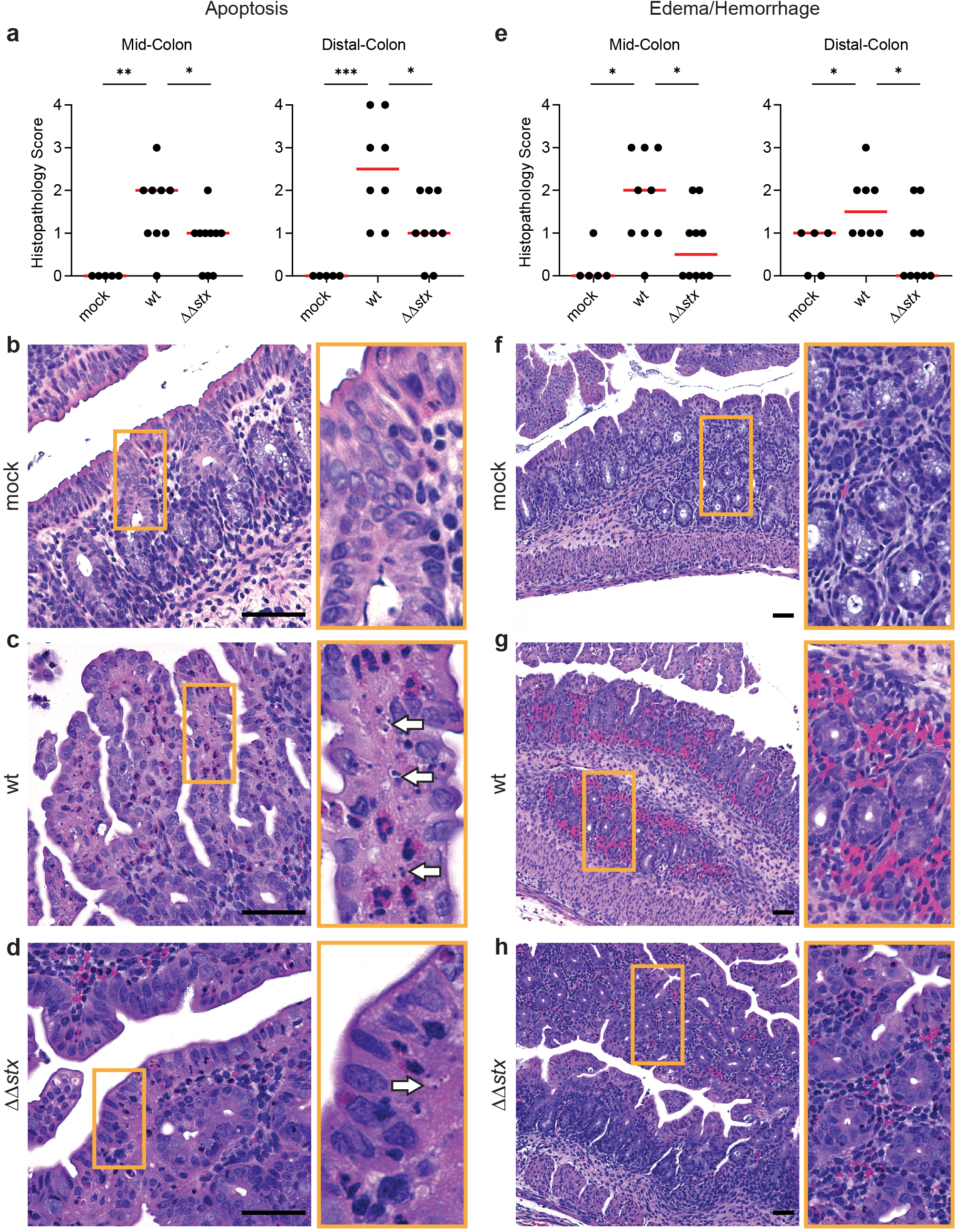
Apoptosis and hemorrhage/edema in the colon is more prominent in animals infected with WT vs ΔΔ*stx* EHEC. (a) Apoptotic nuclei in colon sections from infant rabbits inoculated with PBS (mock), WT or ΔΔ*stx* EHEC 36 hours post inoculation. Scores for individual tissue sections are plotted with the median (red line). Statistical comparisons between groups were made using a two-tailed Mann-Whitney U. P-values were considered significant at less than 0.05 (*), 0.01 (**), or 0.001 (***). (b-d): Example images from mock (score=0), WT (score=4), and ΔΔ*stx* (score=2) infected colons. Scale bars indicate 50 µm. White arrows indicate apoptotic nuclei. Orange box denotes inset displayed to the right. (e): Severity of hemorrhage/edema. (f-h): Representative example images from mock (score=0), WT (score=3) and ΔΔ*stx* (score=1) infected colons. Scale bars indicate 50 µm. Orange box denotes inset displayed to the right.

### Shiga toxin shapes the colonic mucosal transcriptomic response to EHEC

To further investigate how Stx modifies the colonic mucosa’s response to EHEC infection, we used RNA-seq to characterize the transcriptomes of colonic epithelial and lamina propria cells derived from infant rabbits orogastrically inoculated with WT, ΔΔ*stx*, or PBS (mock). In the colon, epithelial cells make initial contact with EHEC and Stx; beneath the epithelial layer, cells in the lamina propria, which include immune cells, respond to signals from the epithelial cells or potentially from direct contact with PAMPs to trigger additional immune responses (61). To profile transcriptional changes in these cell populations, we harvested colons 36 hours post inoculation and collected epithelial and lamina propria cells from 3 rabbits per inoculum type (62).

RNA was extracted, subjected to next-generation sequencing, and mapped to the rabbit genome. Expression of two marker genes—*EPCAM,* which encodes for an epithelial cell adhesion protein, and *PTPRC,* which encodes the immune cell surface marker CD45—confirmed the enrichment of epithelial and immune cells from the EDTA- and collagenase-treated samples respectively (Fig S4A). Next, the global gene expression profiles for individual samples were evaluated and compared using the differential gene expression package DESeq2 (63). Principal component analysis (PCA) revealed that epithelial samples from each group clustered separately, highlighting the specificity of the transcriptomic responses elicited by each inoculation type in this compartment (Fig 2A). PCA did not segregate the lamina propria samples as neatly based on the presence or absence of *stx* (Fig S4B), suggesting that the transcriptional response to Stx is concentrated in the epithelial compartment.

**Figure 2:**
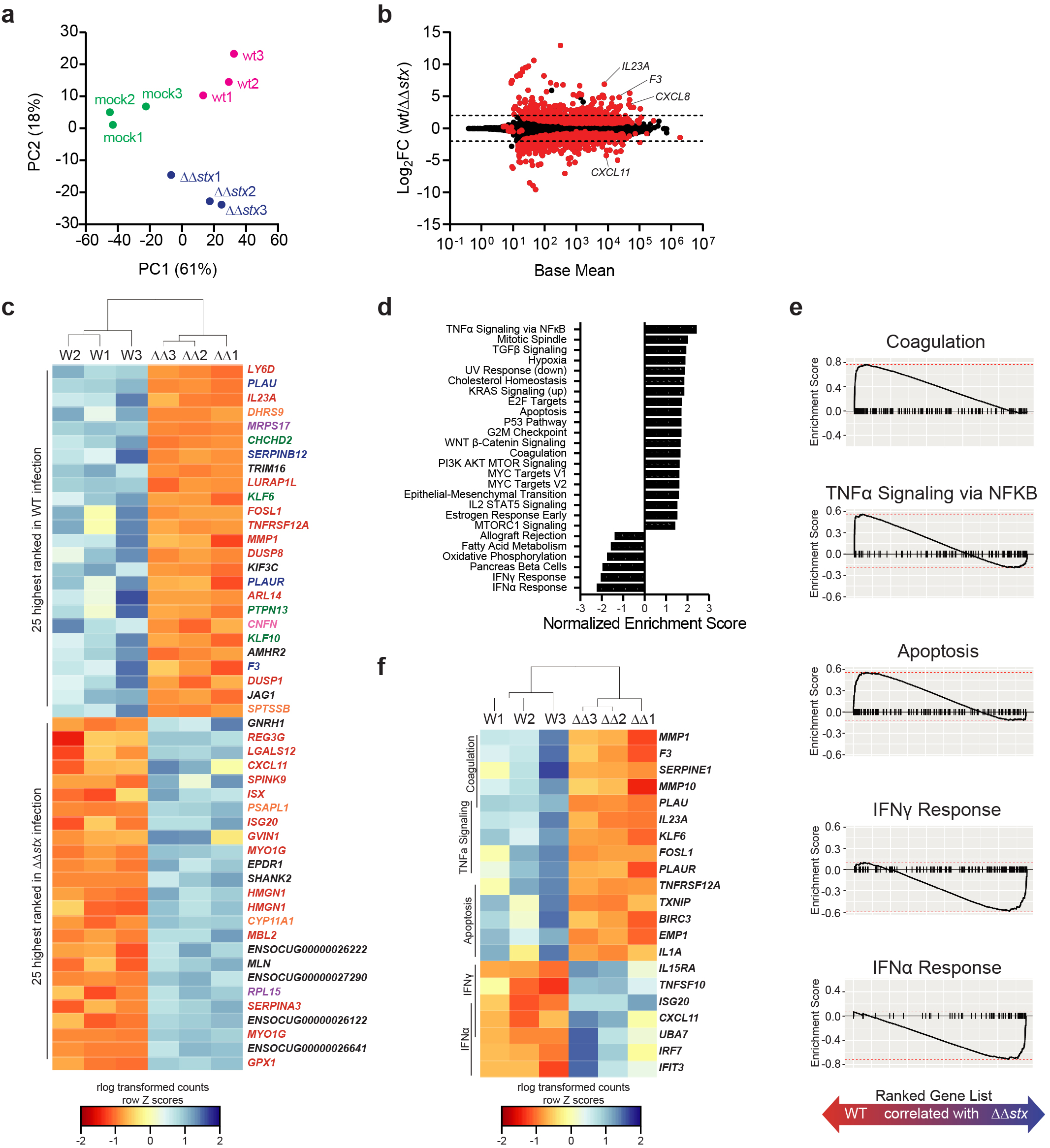
Profiles of colonic epithelial cell transcriptional responses differ between animals infected with WT and ΔΔStx EHEC. a) Principal Component Analysis (PCA) of rlog-transformed expression values of epithelial cells b) Average expression level (base mean) and log2 fold change of transcript abundance in colonic epithelial cells from rabbits inoculated with WT or ΔΔ*stx* EHEC from DESeq2. Genes with significantly different (adjusted p-value < 0.05) transcript abundance are highlighted in red. Dashed line indicates log2 fold change of 2 and -2. c) Heat map of rlog-transformed read counts for 3 animal replicates (WT or ΔΔ*stx* EHEC infected) for top 25 and bottom 25 genes by rank. Hierarchical clustering performed using Euclidian sample distances. Rows are normalized by Z-score. Gene names are colored by function: red – immune, orange – metabolism, green – proliferation/apoptosis, blue – coagulation, purple – translation/protein folding, pink – barrier function/cytoskeleton, black – uncharacterized or other. d) Gene sets from the MiSigDB Hallmark Gene Set Collection significantly associated with WT infection (enrichment score >1) or ΔΔ*stx* infection (enrichment score <-1). e) Gene set enrichment plot for selected pathways. Black tick marks represent genes within pathway organized by rank. f) Heat map rlog-transformed read counts for top 5 genes in the leading edge for indicated selected pathway. Genes organized by rank. Hierarchical clustering performed using Euclidian sample distances

Relative to the mock infected samples, WT infection stimulated more transcriptomic changes than ΔΔ*stx* infection in colonic epithelial cells. Of the ∼30,000 rabbit genes surveyed, 1126 had significantly different expression in WT/mock; for ΔΔ*stx*/mock, 497 genes were differentially expressed (Fig S5) (Dataset S1, S2). 321 genes were in common between the two infection types (Fig S6A). Most of these shared genes were associated with GO Molecular Function terms commonly linked to bacterial infection, including cytokine/chemokine signaling and binding, and LPS recognition through TLR4 signaling (Fig S6B), implying that many typical pathogen-response pathways are activated similarly in the presence or absence of Stx. Similar patterns were observed in the lamina propria transcriptome (Fig S6CD, Dataset S3, S4).

Although the WT and ΔΔ*stx* strains stimulated a shared set of transcripts, we also found robust and widespread differences in the epithelial transcriptomic responses to WT vs ΔΔ*stx* infection. 390 genes exhibited significantly different expression between these two conditions (Fig 2B, Dataset S5). Genes were ranked by adjusted p-value, and hierarchical clustering was performed on rlog transformed read counts. The clustering analysis confirmed that WT and ΔΔ*stx* infection elicit markedly distinct transcriptomic signatures in the epithelium (Fig S4C). Many of the top 50 genes by rank were involved in processes related to coagulation and immune signaling (Fig 2C). Gene set enrichment analysis (GSEA) was performed using fGSEA (64) to further identify transcriptional processes associated with WT or ΔΔ*stx* infection. Using the Hallmark pathways, a highly curated collection of 50 gene sets that describe a wide variety of biological states (65), we identified a number of pathways specifically associated with WT or ΔΔ*stx* infection (Fig 2D, Dataset S6). Notably, apoptosis, coagulation, and NFKB signaling were associated with the transcriptomic response to WT infection, whereas pathways linked with IFNα and IFNγ signaling were associated with ΔΔ*stx* infection (Fig 2E). Using hierarchical clustering on rlog transformed counts of the top five genes from each enrichment-driving leading-edge subset (66), we observed dramatic differences between the transcriptional responses to WT and ΔΔ*stx* infection (Fig 2F). Specifically, the drivers of coagulation, including the gene coding for tissue factor *F3,* as well as pro-inflammatory cytokines *IL23A* and *IL1A* were specifically associated with WT infection (Fig 2F). In ΔΔ*stx* infection, many interferon-stimulated genes (ISGs) were differentially upregulated, including the T-cell chemokine *CXCL11.* Collectively, these analyses suggest that Stx skews epithelial cell innate immune response to EHEC.

In samples from the lamina propria, far fewer genes exhibited differential expression in comparisons between WT and ΔΔ*stx*-infections than in epithelial samples (91 vs 390) (Fig 3A, Dataset S7). Similar to the PCA analysis (Fig S4B), hierarchical clustering did not separate the transcriptional profiles of WT and ΔΔ*stx*-infected colons as clearly as observed in epithelial samples (Fig S4D vs Fig 2A). However, clustering analysis showed that the top 50 genes by rank were distinguishable by inoculum type (Fig 3B). Many of these genes function in coagulation and immune signaling pathways, including several matrix metalloproteases (MMPS), immune signaling pathways including IFNγ, ISGs, and *SLAMF6* and *CD207* (Fig 3C). GSEA with the hallmark pathways confirmed that IFN signaling was associated with ΔΔ*stx* infection (Dataset S8).

**Figure 3:**
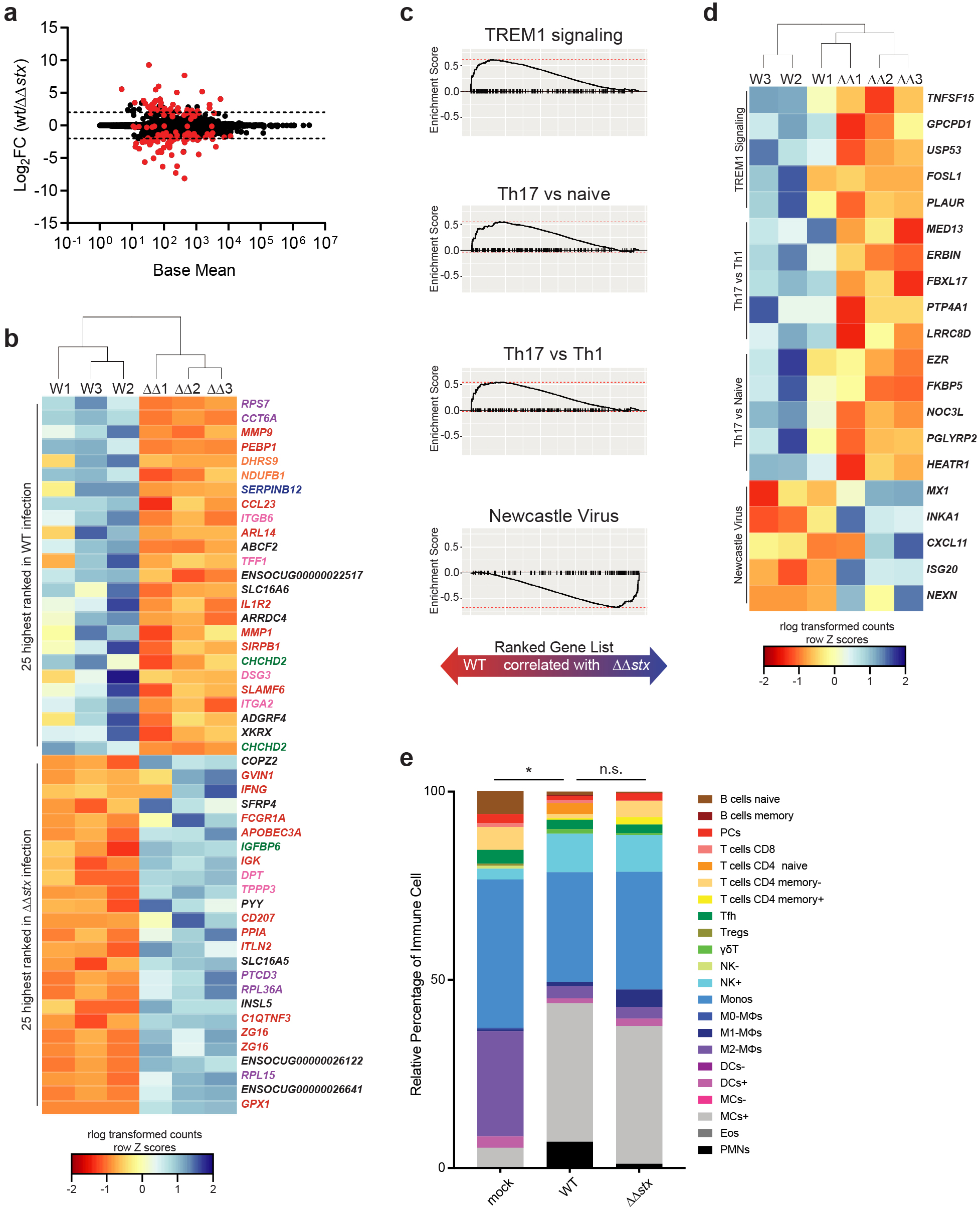
Profiles of colonic lamina propria cell transcriptional responses differ between animals infected with WT and ΔΔStx EHEC. a) Average expression level (base mean) and log2 fold change of transcript abundance in colonic lamina propria cells from rabbits inoculated with WT or ΔΔ*stx* EHEC. Genes with significantly different (adjusted p-value < 0.05) transcript abundance are highlighted in red. Dashed line indicates log2 fold change of 2 and -2. b) Heat map of rlog-transformed read counts for 3 animal replicates (WT or ΔΔ*stx* EHEC infected) for top 25 and bottom 25 genes by rank. Hierarchical clustering performed using Euclidian sample distances. Rows are normalized by Z-score. Gene names are colored by function: red – immune, orange – metabolism, green – proliferation/apoptosis, blue – coagulation, purple – translation/protein folding, pink – barrier function/cytoskeleton, black – uncharacterized or other. c) Gene set enrichment plot for selected pathways from the ImmSigDB Hallmark Gene Set Collection significantly associated with WT infection (enrichment score >1) or ΔΔ*stx* infection (enrichment score <-1). Black tick marks represent genes within pathway organized by rank. d) Heat map rlog-transformed read counts for top 5 genes in the leading edge for indicated selected pathway. Genes organized by rank. Hierarchical clustering performed using Euclidian sample distances. f) Relative cell abundance of immune cell subtypes determined by CIBERSORTx bulk RNA-seq deconvolution. Distributions compared with the Kolmogorov-Sokolov test, p<0.05 (*).

GSEA was also carried out using ImmuneSigDB, a collection of nearly 5,000 gene-sets related to immune system perturbations and immune cell states (Dataset S8) (67). Pathways of interest associated with WT infection included “Th17 vs Th1,” “Th17 vs Naive,” and “TREM1 signaling” revealing that signatures associated with type 3 immunity are present in the lamina propria (Fig 3C). Gene-sets associated with ΔΔ*stx* infection included “Newcastle Virus” which induces a strong interferon response (Fig 3C). Examining the leading-edge subset of these pathways underscored that genes associated with Th17 cells are associated with WT infection while IFNγ-stimulated genes such as *CXCL11* are associated with ΔΔ*stx* infection in the lamina propria (Fig 3D). Together, these analyses suggest that Stx biases lamina propria transcriptional responses toward expression of genes associated with type 3 immunity and that in the absence of this toxin, IFN-related pathways are more prominent.

Enrichment of differentially expressed genes representative of particular pathways could in part reflect infiltration of cells into a tissue in addition to alterations in expression patterns of resident cells. Both WT and ΔΔ*stx* infection led to similar and significant immune cell infiltration in the colon (Fig S2). We attempted to characterize these infiltrates further leveraging the bulk RNAseq data and the deconvolution algorithm CIBERSORTx (68, 69), which relies on cell-type signature matrices to define cell types of interest, to reveal the relative proportion of cell types in each sample. CIBERSORTx and LM22, a matrix of 22 immune cell types that has been successfully applied to define cell types from bulk RNAseq data and has been extensively validated (68, 70), was used to analyze the relative proportion of immune cell types in the lamina propria samples from WT, ΔΔ*stx* and control animals. The relative proportion of immune cell types in the WT infected animals was significantly different than the uninfected (mock) samples, but not different than the distribution in the ΔΔ*stx* infected animals (Fig 3F). The infected tissue sections were associated with a higher relative proportion of activated NK cells, activated mast cells, and neutrophils and lower proportion of M2 macrophages compared to uninfected tissue (Fig 3F, FigS4E). Thus, as in the histologic sections (Fig S2), these analyses suggest that both WT and ΔΔ*stx* stimulate similar immune cell infiltration. Therefore, the differences in the transcriptional profiles observed in animals infected with these strains are likely attributable to Stx-induced changes in the transcriptional landscape of cells resident in the mucosa, rather than differential immune cell infiltration.

### EHEC stimulates expression of coagulation-associated genes in a Stx-dependent manner

Comparison of expression profiles from epithelial and lamina propria samples from animals infected with WT vs ΔΔ*stx* EHEC revealed differences in many genes associated with coagulation. Samples from WT infection had higher levels of transcripts for *F3,* a gene which encodes the initiator of the clotting cascade, MMPs and SERPINs, which are proteases that regulate the processing of coagulation cascade proteins, as well as urokinase and the urokinase receptor, which regulate fibrin deposition. Stx is known to cause thrombosis in vascular beds outside of the GI tract, and has been investigated in the kidney microvasculature (26, 71, 72), but comparatively few studies have focused on Stx-linked coagulation in the intestine.

As antibodies to detect rabbit proteins in tissue are not readily available, we used RNA FISH (RNAscope) (8) to investigate the localization of transcripts of interest identified in the RNAseq data. We probed for Tissue Factor (*F3*) transcripts in infected and control samples and found that there was markedly greater *F3* expression in samples from WT-infected vs ΔΔ*stx*-infected or control rabbits (Fig 4A). *F3* transcripts were observed in >10% of total DAPI+ colon tissue in WT infection and in only ∼0.1% of tissue in ΔΔ*stx* infected animals (Fig 4B). Mean fluorescence intensity (MFI), a proxy for transcript abundance, was much higher in WT vs ΔΔ*stx* samples and the MFI in the latter samples did not differ from the values found in mock infected samples (Fig 4C), suggesting that Stx is critical for stimulating *F3* expression. Notably, almost all (>90%) of the F3-hybridizing signal in WT samples was detected in cells expressing E-Cadherin, an epithelial marker (Fig 4DE). Thus, Stx, when present, appears to primarily act on the colonic epithelium to stimulate *F3* expression. The uniformity of *F3* expression along the epithelium, even in sections where few EHEC cells were detected (Fig S7), suggests that Stx may diffuse along the epithelium to modify transcriptional programs in cells that do not have attached EHEC; alternatively, epithelial cells with attached EHEC may secrete factors that modify transcription in neighboring cells.

**Figure 4:**
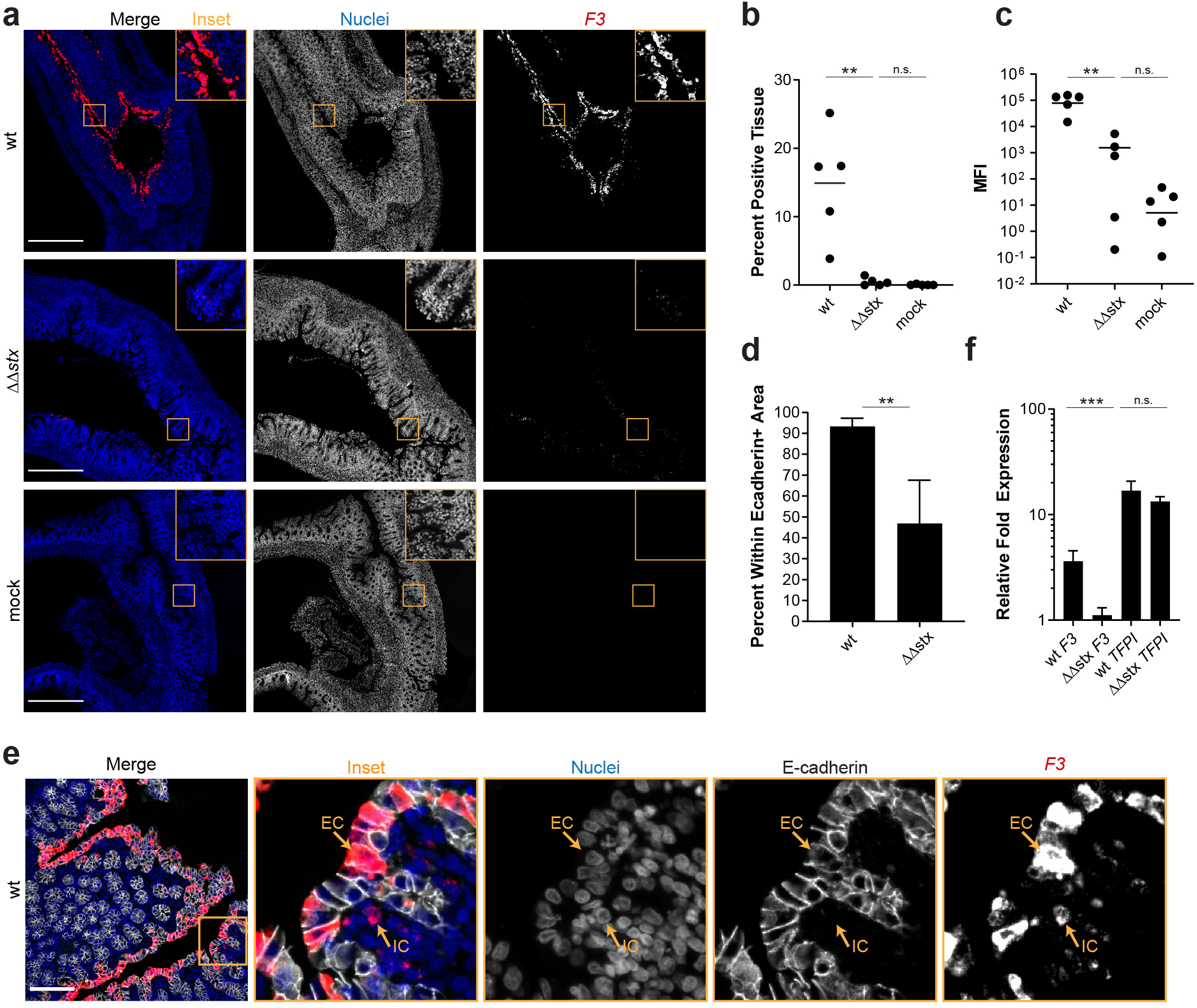
Expression of *F3* in epithelial cells is much greater in animals infected with WT vs ΔΔ*stx* EHEC. a) Immunofluorescence micrographs of colon sections from rabbits inoculated with WT EHEC, ΔΔ*stx* EHEC, or PBS (mock) stained with an RNAscope probe for rabbit *F3* mRNA (red) and DAPI (blue). Scale bar is 500 µM. b) Percentage of tissue section with *F3* signal from individual rabbit colons. Distributions compared using the Mann-Whitney U test, p<0.01 (**), n.s. indicates not significant. c) Quantification of mean fluorescent intensity (MFI) of pixels above background from individual rabbit colons plotted with mean. Distributions compared using the Mann-Whitney U test, p<0.01 (**), n.s. indicates not significant. d) Percent *F3* signal within E-cadherin positive cells. Distributions compared using the Mann-Whitney U test, p<0.05 (*). e) Immunofluorescence micrographs of sections stained with an RNAscope probe to rabbit *F3* mRNA (red), DAPI (blue), and anti-E-cadherin antibody (white). Scale bar is 500 µM. IC indicates immune cell, EC indicates epithelial cell. f) Normalized expression of *F3* and *TFPI* measured by RT-qPCR in HT29 colonic epithelial cells exposed to WT EHEC, ΔΔ*stx* EHEC, or PBS for 6 hours. Expression was normalized to the housekeeping gene *GAPDH* and then expressed relative to basal expression in uninfected cells. Statistical significance determined by a Students two-tailed t-test, p < 0.001 (***), n.s. indicates not significant.

We also explored whether Stx stimulates *F3* transcription in HT29 cells, a human colonic epithelial cell line (74). qPCR was used to quantify *F3* transcripts in HT29 cells after infection with WT, ΔΔ*stx* EHEC, or PBS (mock). With WT infection, *F3* gene expression was 5-fold higher than in uninfected cells (Fig 4F); moreover, as in rabbits, induction of *F3* expression in HT29 cells was largely dependent on Stx and there was little difference in *F3* expression in uninfected cells vs those infected with ΔΔ*stx* infection (Fig 4F). It has been reported that Stx leads to increases in Tissue Factor pro-coagulant protein activity in renal proximal tubule cells and endothelial cells (75–78); this effect is thought to be primarily driven by a decrease in the expression of Tissue Factor Protein Inhibitor (TFPI), and not an increase in *F3* gene expression (75, 79). We also measured *TFPI* transcript levels in the samples used to measure *F3* expression, to assess if this pathway is also active in HT29 cells. Both WT and ΔΔ*stx* infection similarly stimulated levels of *TFPI* mRNA >10-fold compared to the uninfected cells (Fig 4F). Thus, Stx appears to regulate Tissue Factor by different mechanisms in colonic epithelial cells vs endothelial cells.

### Stx biases colonic immune responses toward type 3 immunity

Analyses presented above revealed that Stx skews colonic mucosal gene expression to EHEC infection, stimulating expression of several pro-inflammatory cytokines, such as IL23, relative to levels observed in ΔΔ*stx* infection (Fig 2, 3). We used RNA FISH to compare the fluorescence intensity and distribution of *IL23a* transcripts in colons from rabbits infected with WT or ΔΔ*stx* (Fig 5A). The intensity and percent of tissue expressing *IL23A* signal in samples from animals infected with WT EHEC were markedly higher than those in animals infected with ΔΔ*stx* EHEC (Fig 5BC). These values did not differ in ΔΔ*stx* and control samples, suggesting that Stx stimulates *IL23A* expression. Unexpectedly, nearly all of the *IL23A* expression in WT samples was detected within epithelial cells (E-cadherin positive cells) compared to cells in the lamina propria (Fig 5D).

**Figure 5:**
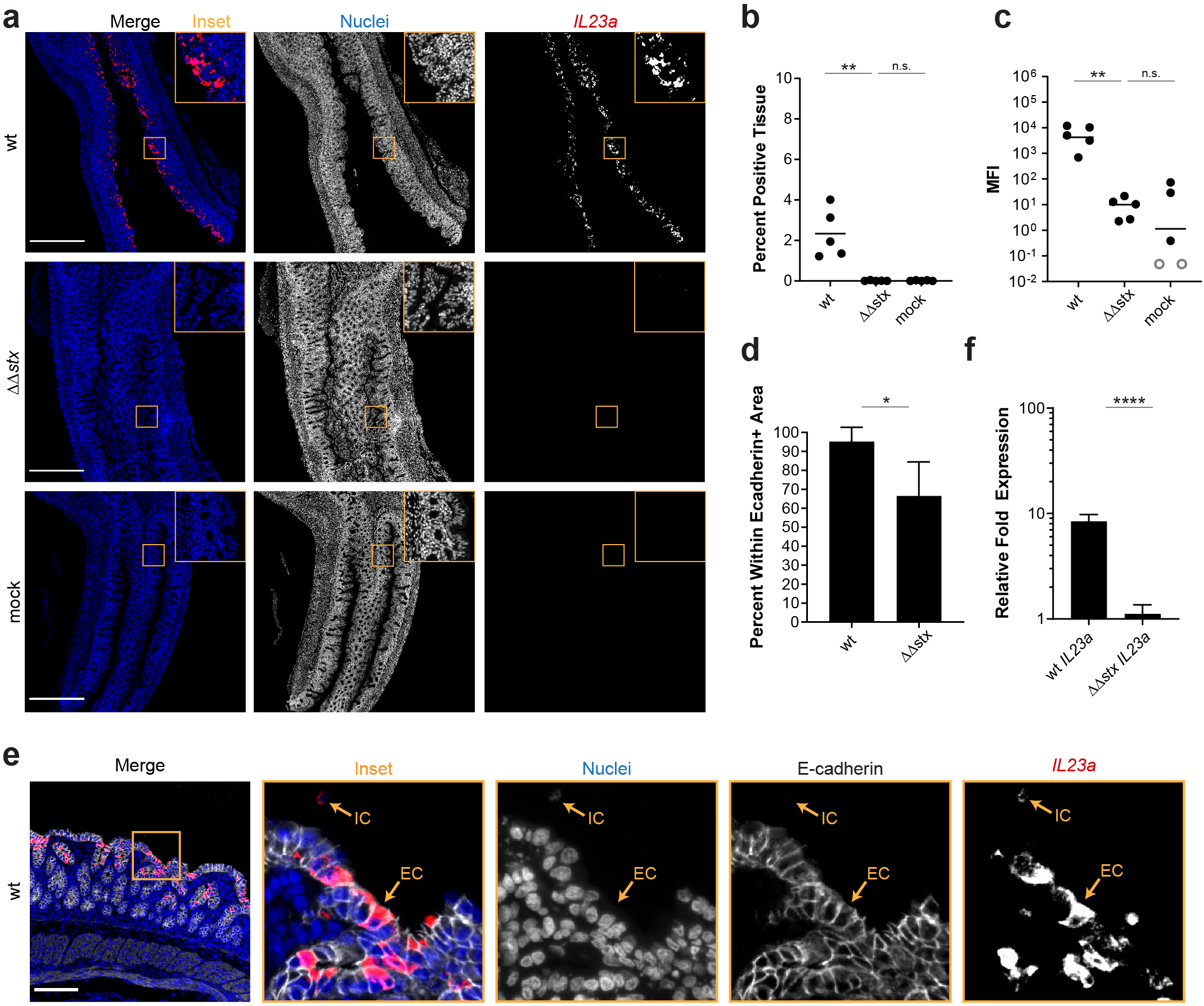
Expression of *IL23a* in epithelial cells is much greater in animals infected with WT vs ΔΔ*stx* EHEC. a) Immunofluorescence micrographs of colon sections from rabbits inoculated with WT EHEC, ΔΔ*stx* EHEC, or PBS (mock) stained with an RNAscope probe for rabbit *IL23a* mRNA (red) and DAPI (blue). Scale bar is 500 µM. b) Percentage of tissue section with *IL23a* signal from individual rabbit colons. Distributions compared using the Mann-Whitney U test, p<0.01 (**), n.s. indicates not significant. c) Quantification of mean fluorescent intensity (MFI) of pixels above background from individual rabbit colons plotted with mean. Distributions compared using the Mann-Whitney U test, p<0.01 (**), n.s. indicates not significant Open circles indicate limit of detection. d) Percent *IL23a* signal within E-cadherin positive cells. Distributions compared using the Mann-Whitney U test, p<0.05 (*). e) Immunofluorescence micrographs of sections stained with an RNAscope probe to rabbit *IL23a* mRNA (red), DAPI (blue), and anti-E-cadherin antibody (white). Scale bar is 500 µM. IC indicates immune cell, EC indicates epithelial cell. f) Normalized expression of *IL23a* measured by RT-qPCR in HT29 colonic epithelial cells exposed to WT EHEC, ΔΔ*stx* EHEC, or PBS for 6 hours. Expression was normalized to the housekeeping gene *GAPDH* and then expressed relative to basal expression in uninfected cells. Statistical significance determined by a Students two-tailed t-test, p < 0.001 (***), n.s. indicates not significant.

IL23 also promotes the expression of other cytokines such as the chemokine CXCL8, which is a neutrophil chemoattractant. Similar to *F3* and *IL23A* expression, *CXCL8* transcripts were observed at much greater intensity and in a much larger area of tissue in samples from WT vs ΔΔ*stx* infection (Fig S8). Furthermore, these *CXCL8* transcripts were primarily present in epithelial cells (Fig S8). Similar to *F3, CXCL8* signal was uniform throughout epithelial tissue, showing no apparent correlation with EHEC foci along the epithelium (Fig S7). Together these observations suggest that, during EHEC infection, Stx biases the host immune response toward type 3 immunity by stimulating IL23 expression in the colonic epithelium.

In the RNAseq data, we found that animals infected with ΔΔ*stx* had statistically significant higher levels of *IFNG* transcripts than those infected with WT (Fig 3B), suggesting that Stx may inhibit production of IFNγ, a cytokine linked to type 1 immunity. Reduction in IFNγ-mediated STAT-1 phosphorylation has been linked to Stx activity in vitro (80). In vivo, IFNγ is produced mainly by Th1 cells, ILC1s, NK cells and macrophages in the lamina propria. IFNγ signaling stimulates transcription of many other genes known as interferon-stimulated genes (ISGs) (81), many of which were found to be differentially expressed in animals infected with WT vs ΔΔ*stx* (Fig 2, 3). One of those genes, *CXCL11,* is an IFNγ-inducible chemokine produced by macrophages that acts as a chemoattractant for CXCR3+ T cells; macrophage engagement of CXCR3+ T cells promotes Th1 cell development. RNA FISH analyses revealed that the MFI of the *CXCL11* signal was more intense and detected in more tissue area in samples from animals infected with ΔΔ*stx* vs WT (Fig 6ABC). Interestingly, approximately 50% of the *CXCL11* transcript signal was found in epithelial cells (Fig 6DE). These observations suggest that Stx inhibits IFNγ signaling in the colonic epithelium during EHEC infection, using this cellular layer to tip the host response towards type 3 immunity.

**Figure 6:**
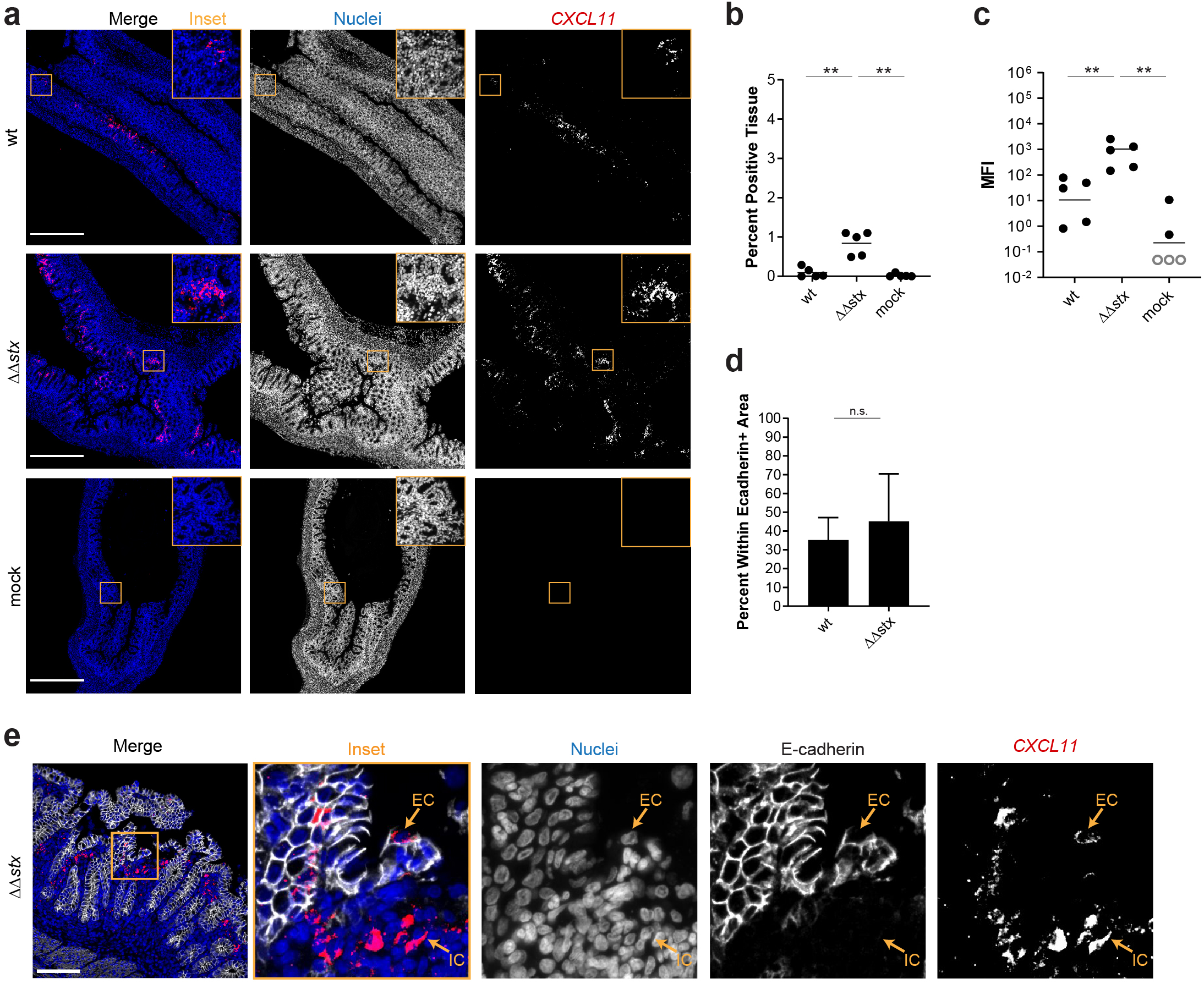
Expression of *CXCL11* in is greater in animals infected with ΔΔ*stx* vs WT EHEC. a) Immunofluorescence micrographs of colon sections from rabbits inoculated with WT EHEC, ΔΔ*stx* EHEC, or PBS (mock) stained with an RNAscope probe for rabbit *CXCL11* mRNA (red) and DAPI (blue). Scale bar is 500 µM. b) Percentage of tissue section with *IL23a* signal from individual rabbit colons. Distributions compared using the Mann-Whitney U test, p<0.01 (**), n.s. indicates not significant. c) Quantification of mean fluorescent intensity (MFI) of pixels above background from individual rabbit colons plotted with mean. Distributions compared using the Mann-Whitney U test, p<0.01 (**), n.s. indicates not significant. Open circles indicate limit of detection. d) Percent *CXCL11* signal within E-cadherin positive cells. Distributions compared using the Mann-Whitney U test, p<0.05 (*). e) Immunofluorescence micrographs of sections stained with an RNAscope probe to rabbit *CXCL11* mRNA (red), DAPI (blue), and anti-E-cadherin antibody (white). Scale bar is 500 µM. IC indicates immune cell, EC indicates epithelial cell.

## Discussion

Orogastric inoculation of infant rabbits with EHEC leads to diarrheal disease and colonic pathology that closely resembles many of the intestinal aspects of human EHEC infection (28, 30, 58). Here, we used this model to study how Stx modifies the host response to infection in the colon by comparing the histopathology and transcriptional profiles of the colonic mucosa from animals infected with WT EHEC or an isogenic mutant lacking Stx genes (ΔΔ*stx*). We found that Stx, a potent toxin, increases apoptosis and hemorrhage in colonic tissue and dramatically remodels the colonic epithelium’s transcriptional response to EHEC infection. Though the transcriptional responses elicited by WT and ΔΔ*stx* EHEC infection exhibited some overlap, particularly in pathways commonly associated with gram-negative infection, these isogenic strains provoked very distinct transcriptional profiles in the colonic epithelium. Nearly twice as many genes were differentially regulated in animals infected with WT/mock vs ΔΔ*stx*/mock, and many of the differences in the epithelial response related to immune signaling and coagulation. Overall, Stx appears to skew the epithelial transcriptional response toward expression of genes associated with type 3 immunity.

The cellular response to Stx has also been characterized using tissue-cultured cells. The ‘ribotoxic stress response’ to Stx-mediated cellular damage leads to the upregulation of a variety of genes and production of proteins which modulate the immune response (23, 25, 26). Purified Stx promotes the production of transcription factors (including JUN and FOS) and inflammatory cytokines (CCL2, CCL3, CCL4, CCL5, CSF2, CSF3, CXCL1, CXCL2, CXCL3, CXCL5, CXCL8, IL10, IL1αβ, IL6, TNFα) in cultured epithelial and immune cells (42, 41, 43, 22, 40, 29, 44–47, 26). We found that several of these genes, including *JUN, FOS, CCL4, CXCL8,* and *IL1A* were differentially expressed in WT vs ΔΔ*stx* infected colons; thus, Stx by itself, in the absence of additional EHEC-derived factors may account for a subset of the transcriptional changes we identified in the colonic epithelium of animals infected with the WT strain. However, recent transcriptomic studies of the response of human intestinal organoids did not detect differential expression of many of the coagulation-associated or immune signaling genes (including *F3, CXCL8*, and *IL23A*) that we found in infected infant rabbits (47, 51); these differences are potentially explained by organoid culture conditions, which can limit cytokine expression (82). Overall, these discrepancies highlight the important differences in gene expression patterns observed in cultured cells and the highly complex milieu of the host intestine, and that use of purified toxin is not sufficient to capture the intricacies of the host response to a pathogen with a variety of immunomodulating signals such as LPS and T3SS effectors.

Comparisons of the epithelial transcriptional profiles induced by WT and ΔΔ*stx* EHEC in rabbit colons suggest that Stx promotes the transcription of genes related to type 3 immunity. The expression of *IL23A,* the subunit p19 of the type 3 cytokine IL23, was markedly higher in WT than ΔΔ*stx* infected colons. IL23 drives the immune response towards type 3 immunity, which is often activated in response to extracellular pathogens (54), and is generally thought to be produced by dendritic cells (83). Unexpectedly, nearly all of the *IL23A* signal was detected in colonic epithelial cells. However, we did not detect epithelial transcripts for *IL12B*, which encodes for p40, the other subunit of the IL23 protein. Typically, the two subunits are expressed together and form the IL23 heterodimer (84), but the gene expression of these two subunits is not always temporally synchronized (85). It is possible that IL23 p19 could have an independent function or an alternative partner in intestinal epithelial cells, as has been suggested recently (86). In contrast to the induction of *IL23A,* Stx appears to inhibit expression of *IFNG* and many ISGs such as *CXCL11,* since these cytokines and downstream factors were downregulated in WT compared to ΔΔ*stx* infected colons. IFNγ is the characteristic cytokine of type 1 immunity and is typically activated to protect cells against intracellular microbes (54). In vitro studies have also shown that Stx can suppress IFNγ-mediated signaling (80), while similar attaching-and-effacing pathogens which lack Stx do not suppress IFNγ (EPEC) (73) and robustly activate type 1 responses (*Citrobacter rodentium*) (55–57). Additional bacterial toxins with distinct mechanisms of action have also been reported to bias the innate response along this axis: *Clostridioides difficile* toxin and the heat-labile enterotoxin of *E. coli* and have also been shown to promote type 3 immunity by inducing IL23 production (87, 88). Our findings suggest that Stx shapes the early innate immune response to infection, and thus likely has implications for the development of adaptive immune responses and the host’s capacity to clear the infection. Though reagents to verify changes in the abundance of proteins are not readily available for rabbits, we hope that this will be possible in the future.

Stx-mediated damage to endothelial cells, particularly in the renal microvasculature, is a well-studied hallmark of HUS. Toxin damage to endothelial cells triggers the coagulation cascade, which leads to thrombosis with fibrin deposition and hemolysis of RBCs (27). In the colon, patient biopsies have revealed that EHEC infection can also induce microvascular thrombi and fibrin deposition (8), but few studies have investigated the patterns of gene expression which may contribute to thrombosis in the intestine. Although we were unable to detect fibrin deposition in tissue sections, we found that *F3*, the gene encoding the initiator of the coagulation cascade, is dramatically induced in colonic epithelial cells in WT but not ΔΔ*stx* infection. Tissue Factor is typically expressed on cells which are not in contact with blood, such as epithelial cells (89, 90), and can be induced in response to inflammatory stimuli (91). In endothelial cells, Shiga toxin can promote Tissue Factor activity through a mechanism associated with a decrease in expression of Tissue Factor Protein Inhibitor (TFPI) (75–79). However, in rabbit colonic epithelial cells and in the human colon cancer cell line HT-29, *TFPI* gene expression was not altered despite a marked increase in Tissue Factor expression, suggesting an alternate mechanism of Tissue Factor induction in colonic epithelial cells.

Collectively, our findings reveal that Stx powerfully shapes the host response to EHEC. Though the degree of epithelial transcriptional remodeling by Stx is striking, it is not immediately apparent what benefit this program yields for the pathogen, especially with no discernible difference in the burden of WT and ΔΔ*stx* EHEC in infected animals. It is possible that the shaping of the inflammatory response by Stx augments diarrhea and thereby enhances pathogen dissemination. Additionally, altered gene expression patterns during EHEC infection related to coagulation and inflammation are suggestive of ‘thromboinflammation,’ a mechanism in which thrombosis and inflammation synergize to contribute to disease pathology (92). Our findings illustrate the potency of combining isogenic mutants with cellular compartment-specific characterization of host responses to infection, for unravelling how individual virulence factors contribute to the cell type-specific pathogen-host dialogue in disease. For EHEC infection, understanding this dialogue and the innate immune processes contributing to the early colonic phase of disease prior to HUS could offer valuable clues for developing new therapies.

## Materials and Methods

### Ethics Statement

Animal experiments were conducted using protocols approved by Brigham and Women’s Hospital Committee on Animals (Institutional Animal Care and Use Committee protocol number 2016N000334 and Animal Welfare Assurance of Compliance number A4752-01) and in accordance with recommendations in the National Institute of Health’s Guide for the Care and Use of Laboratory Animals and the Animal Welfare Act of the United States Department of Agriculture.

### Bacterial Strains and Growth Conditions

Bacterial strains were cultured in LB medium or on LB agar plates at 37°C. A gentamicin-resistant mutant of *E. coli* O157:H7 strain EDL933 (Δ*lacI*::*aacC1*) (93) was used in all experiments in this study and gentamicin (Gm) was used at 10 μg/mL. The ΔΔ*stx* mutant was constructed using lambda red recombineering (94) as described (95).

### Infant rabbit infection and tissue processing

Two-day old litters of mixed gender New Zealand White rabbits were co-housed with a lactating dam (Charles River). Infection inocula were prepared by diluting 100 μl of overnight culture into 100 mL of LB Gm; then, following 3 hours of growth at 37°C with shaking, 30 units of culture at OD_600_ = 1 (about 8 mL) were pelleted and resuspended in 10 mL PBS. Dilutions of the inoculum were plated to enumerate CFU. Each infant rabbit was orogastrically inoculated with 500 μl of the inoculum (∼1×10^9^ CFU), using a size 4 French catheter. Following inoculation, the infant rabbits were monitored at least 2x/day for signs of illness and euthanized 2 days (36-40 hours) post infection, when the entire intestinal tract was removed.

One cm sections of the medial and distal colon were removed post necropsy and the tissue pieces were homogenized in 1 mL of sterile PBS using a minibeadbeater-16 (BioSpec Products, Inc.). Dilution series of the homogenates were plated on LB Gm plates, which were incubated overnight at 37°C, to determine CFU/g bacterial burdens in tissue sections.

### Tissue preservation and histopathology

Two cm sections of medial and distal colon were fixed in 2 mL 10% neutral-buffered formalin overnight (∼16 hours) at room temperature. The next day, tissue sections were transferred to 2 mL 70% ethanol. Formalin-fixed, paraffin embedded 5 uM sections were stained with hematoxylin and eosin (H&E) by the Rodent Histopathology Core at Dana Farber Cancer Institute. Slides were blindly evaluated by a histopathologist and scored semi-quantitatively. Sections were evaluated for inflammation using the following criteria: 0, none; 1, mild infiltration of immune cells into lamina propria; 2, moderate infiltration; 3, extensive infiltration; 4, severe and extensive infiltration. Apoptosis was evaluated using the following criteria: 0, none; 1, few cells observed with fragmented nuclei; 2, many cells with fragmented nuclei; 3, significant apoptotic nuclei and penetration to crypts; 4, transmural apoptosis. Edema, congestion and hemorrhage were evaluated using the following criteria: 0, none; 1, mild vascular congestion and/or mild edema; 2, moderate congestion and/or edema; 3, congestion with hemorrhage +/ edema; 4, congestion with severe multifocal hemorrhage +/- edema. Heterophil infiltration was evaluated using the following criteria: 0, none; 1, scattered individual heterophils or small clusters in the lamina propria; 2, multifocal aggregates in mucosa with few cells in lumen; 3, multifocal aggregates in mucosa with abundant cell extrusion into lumen; 4, multifocal aggregates in mucosa with large heterophilic intraluminal rafts. Sloughing was evaluated using the following criteria: 0, none; 1, few epithelial cells sloughed from luminal surface; 2, moderate number of epithelial cells sloughed from luminal surface; 3, epithelial surface is severely disrupted; 4, extensive and severe sloughing (epithelial layer is absent). Scores were compared between infection types using a two-tailed Mann-Whitney U statistical test. The Bejmamini-Hochberg Procedure was used to control for the false discovery rate with multiple comparisons at 20%. P-values were considered significant at less than 0.05 (*), 0.01 (**), and 0.001 (***).

### Tissue preparation for RNA-sequencing

Five cm sections from between the medial and distal colon were harvested and processed immediately post necropsy for RNA sequencing from 3 rabbits inoculated with PBS (mock), WT or ΔΔ*stx* EHEC. Epithelial cell and lamina propria cell fractions were isolated from tissue using a method similar to that described previously (62). First, fat was trimmed from the tissue, and luminal contents were gently pressed out. The tissue was then cut longitudinally and rinsed in 1 mL of wash solution (RPMI 1640, 2% Fetal Bovine Serum (FBS), 10mM HEPES, and 100 ug/mL penicillin-streptomycin). Next, the tissue was rinsed in 40 mL ice-cold Ca/Mg-free HBSS before being transferred to 10 mL of epithelial dissociation solution (HBSS, 100 ug/mL penicillin-streptomycin, 10 mM HEPES, 2% FBS, 10mM EDTA) freshly supplemented with an additional 100 uL of 0.5M EDTA. To remove dying and dead epithelial cells, the tissue was incubated in epithelial dissociation solution 37°C at 125 rpm or 5 minutes, then incubated on ice for 5 minutes, then shaken vigorously 10 times and vortexed for 2 seconds. Supernatants were discarded and the tissue piece was transferred into a new tube of 10 mL of epithelial dissociation solution freshly supplemented with an additional 100 uL of 0.5M EDTA. The solution was brought to room temperature quickly by briefly warming in a 37°C bath, then incubated for 20 minutes at 37°C at 125 rpm centrifugal rotation, then incubated on ice for 5 minutes, shaken vigorously 15 times, and vortexed vigorously for 10 seconds. The supernatant was transferred to a fresh tube and centrifuged at 300xg for five minutes. The cell pellet was resuspended in 2 mL Trizol and the solution was stored at −80°C until RNA extraction.

After epithelial cell dissociation, the remaining tissue piece was transferred to a tube containing 5 mL of enzymatic digestion solution (RPMI 1640, 2% Fetal Bovine Serum (FBS), 10mM HEPES, and 100 ug/mL penicillin-streptomycin, fresh 100 ug/mL Liberase TM, fresh 100 ug/mL DNaseI) and incubated at 37°C with centrifugal rotation at 125 rpm for 30 minutes. The digestion was quenched by adding 80 uL of 0.5M EDTA. The solution was filtered through a 40 uM cell strainer and rinsed with HBSS to a final volume of 30 mL. This tube was spun down at 400xg for 10 minutes, and the cell pellet was resuspended in 2 mL Trizol. Samples were stored at −80°C until RNA extraction.

### RNA extraction and mRNA seq library preparation and sequencing

RNA was extracted from Trizol using the Direct-Zol RNA MiniPrep Plus kit from Zymo with some modifications. First, Trizol samples were incubated at 65°C until just thawed (5-10 minutes). One mL samples were added to RNase-free microcentrifuge tubes and 200 uL of chloroform was added to each tube. The tubes were inverted 10x for mixing, and incubated at room temperature for 3 minutes. The samples were spun at 12,000xg for 15 minutes at 4°C, to separate the aqueous and organic layers. The clear aqueous phase was removed, an equal volume of 100% ethanol was added, and the sample was mixed by inversion ten times before incubating at room temperature for 5 minutes. The entire volume was transferred to a Direct-Zol spin column and spun for one minute. The samples were washed with pre-wash buffer twice, with wash buffer once, spun empty to remove residual buffer twice, and eluted with 50 uL RNase-free water. Total RNA was assessed for quality and integrity (RINe) using a High Sensitivity RNA ScreenTape (Agilent) at the HMS Biopolymers Facility.

RNA of high quality (RINe>8) was prepared for mRNA-sequencing using the KAPA mRNA HyperPrep kit (Roche). Libraries were quantified using a High Sensitivity D1000 ScreenTape (Agilent) and High Sensitivity Qubit. Libraries were sequenced on a NextSeq 550.

### Differential expression analysis

First, resources for the rabbit genome were compiled. We concatenated FASTA files for each rabbit chromosome, mitochondrial DNA, and unplaced scaffolds from OryCun2.0 (Assembly GCA_000003625.1) to create a reference genome FASTA file. The Ensembl annotation (May 2019) was used. Sequencing reads and genome resources were uploaded to the Galaxy web platform (96), and the public server usegalaxy.org was used to process and map reads. First, reads were trimmed using Trim Galore! (Galaxy Tool version 0.4.3.1) with automatic adapter sequence detection. Then, trimmed reads were mapped to the rabbit reference genome and annotation using RNA STAR (Galaxy Tool version 2.6.0b-1). featureCounts (Galaxy Tool version 1.6.4+galaxy1) was used to build a count matrix from mapped reads using the Ensembl annotation as a guide. Count matrices were exported from Galaxy and imported into R (version 3.5.3) (97). The read counts for each gene were normalized to Transcripts Per Kilobase Million (TPM) to compare expression of marker genes across samples. TPM was calculated by first dividing the number of read counts by the length of the gene in kilobases to yield reads per kilobase (RPK). Gene RPKs were summed for each sample and this number was divided by 1,000,000 to yield a sample-specific scaling factor. The RPK value for each gene was divided by the scaling factor to yield TPM. The non-normalized count matrix was also analyzed using DESeq2 (version 1.22.2) (63) to compare the abundance of transcripts between different inoculum types to identify differentially expressed genes. Parametric dispersion was used and shrinkage of effect size was performed using the package apeglm (98). Genes with an adjusted p-value of less than 0.05 were considered to be differentially expressed. Normalized read counts were generated using a regularized log (r-log) transformation and used to perform principal component analysis (PCA) of each sample. Raw reads and count matrices have been deposited into the Gene Expression Omnibus (GEO) repository (accession number pending).

### Hierarchical clustering

For each gene, the parameter ‘rank’ was calculated by multiplying the sign of the fold-change by the log10 transformed adjusted p-value from DESeq2. Genes were ordered by rank, and hierarchical clustering of samples was performed using the gplot (version 3.0.1) function Heatmap.2 using r-log transformed read counts, constructing a dendrogram by column (samples). Heatmaps were scaled by row and colors assigned using ColorBrewer palette RdYlBu (version 1.1.2) (99). Similar clustering analysis was performed for the top 50 genes by rank, and genes in the leading-edge subset from GSEA.

### Gene set enrichment

Gene set enrichment was performed using fast GSEA (fGSEA) in R (version 1.8.0) (64). Only genes with annotation were considered. Genes were ranked with the parameter ‘rank’. The hallmark gene sets (65) and immunological signatures gene sets (67) from MSigDB (100) were used. 1000 permutations were completed, and categories with less than 10 genes and greater than 500 genes were excluded. Pathways were considered to be significantly enriched if the adjusted p-value was less than the false-discovery rate of 5%. Enrichment scores were plotted using the plotEnrichment function in R.

### Comparing differentially expressed genes in infected vs uninfected tissue

Genes with an adjusted p-value of <0.05 and log2 fold-change of >2 or <-2 from DESeq2 (WT vs mock, ΔΔ*stx* vs mock) were considered differentially expressed. These lists were compared using the online tool BioVenn (101). The list of commonly differentially expressed genes from these comparisons was analyzed using the webtool g:Profiler to perform functional enrichment analysis (g:GOst) (102).

### CIBERSORTx

CIBERSORTx (68, 69) was performed using the webtool (https://cibersortx.stanford.edu/) to impute cell fractions. TPM values for all annotated genes were imported as a mixture matrix, and LM22 was used as a signature matrix file. Distributions of the relative abundance of immune cells were compared using a Kolmogorov-Sokolov test. Relative abundance of individual cell types were compared using a Mann-Whitney U test.

### Fluorescent *in situ* RNA hybridization

Fluorescent in-situ hybridization (FISH) was performed using RNAscope (ACDBio) Multiplex Fluorescent V2 Assay in 5 different rabbit colon tissue sections per inoculum type (WT, ΔΔ*stx* and mock). Manufacturer’s protocols were followed and the RNAscope HybEZ oven was used for all incubations. Freshly sectioned formalin-fixed, parrafin embedded (FFPE) tissue sections (5 uM) were processed following manufacturer’s protocol. Briefly, sections were treated with boiling target retrieval buffer for 15 minutes and digested with Protease Plus for 28 minutes. Custom RNAscope C1 probes for rabbit mRNA *IL23a, CXCL8, CXCL11, F3,* or *dapB* (negative control) were hybridized to the tissue. The C1 probe was detected with Opal 570 dye (Akoya Biosciences) diluted 1:1000 in Multiplex TSA buffer (ACDBio). Following completion of the RNAscope assay, sections were processed further for immunofluorescence. Sections were permeabilized with 0.1% TritonX100 for 10 minutes at room temperature, washed, and treated with 5% BSA for 1 hour to block non-specific signal. Sections were washed and stained with primary antibody to E-cadherin (1:100 anti E-cadherin mouse monoclonal antibody, BD Biosciences 610181) and/or primary antibody to O157-antigen (1:2000 anti *E. coli* O157:H7 goat polyclonal antibody, Abcam ab30521) overnight at 4°C. The next day, sections were washed and stained with an anti-mouse secondary antibody conjugated to Alexa 647 and/or anti-goal secondary antibody conjugated to Alexa 488 diluted 1:500 in PBS (Invitrogen A-21235). Sections were then stained with DAPI (2 ug/mL) for 5 minutes before mounting with ProLong Diamond antifade mountant (Thermo Fisher).

### Quantitative image analysis

Samples were imaged with a Nikon Ti Eclipse microscope equipped with a widefield Andor NeoZyla camera and a 20x objective. For H&E stained sections, Kohler alignment and white-balance was performed, and then RGB images were captured. For sections stained with fluorescent antibodies and FISH probes, 4×6 stitched images (10% overlap) were captured for each tissue section using multi-channel acquisition (blue, red, far-red) using 16-bit imaging. In ImageJ/FIJI (103), threshold values for each channel were determined using the sections stained with the negative control probe (*dapB*), which should have no signal in the red channel above background. To analyze mean fluorescence intensity (MFI), the threshold value for the blue channel (DAPI) was set using FIJI to create a binary mask. This mask was applied to the red channel (RNAscope probe), and a histogram of intensity values for pixels within this mask was recorded. The average value above background was recorded for each sample. To analyze the percentage of tissue expressing a transcript of interest, thresholds were applied to both the blue (DAPI) and red (RNAscope) channels to create binary masks. The area of the DAPI mask was recorded. FIJI “create selection” tool was used to draw a selection around the DAPI area. This selection was transferred to the RNAscope channel, and the area of RNAscope signal within this area was recorded. We calculated area of tissue expressing signal by dividing area of RNAscope signal (RNAscope area within DAPI selection) by total tissue area (area of DAPI mask). To calculate the percent of the RNAscope signal derived from epithelial cells, we first applied a threshold to the far-red (E-cadherin) and red (RNAscope) channel to create a binary mask. The area of the RNAscope mask was recorded. Then, the “create selection” tool in FIJI was used to draw a selection around the binary mask of the E-cadherin area. The section was enlarged by 2uM to accommodate signal at the edge of the cells. This section was transferred to the binary masked RNAscope channel. The area of RNAscope binary signal within the E-cadherin selection was recorded. We divided the area of RNAscope signal within E-cadherin selection by the total RNAscope signal to determine percentage of signal within epithelial cells. These three quantifications (mean fluorescent intensity, percent of tissue expressing signal, and percent of signal within epithelial cells) were compared for WT infected vs mock and ΔΔ*stx* infected vs mock for each probe using a two-tailed Mann-Whitney U statistical test. The Bejmamini-Hochberg Procedure was used to control for the false discovery rate with multiple comparisons at 20%. P-values were considered significant at less than 0.05 (*), 0.01 (**), and 0.001 (***).

### Tissue culture infection and RT-qPCR

Human colon colorectal adenocarcinoma cells (HT-29, ATCC HTB-38) were purchased from ATCC and cultured in McCoy’s 5A Medium supplemented with 10% fetal bovine serum. Cells were grown at 37°C with 5% CO2. Two days before infection, 500,000 cells were seeded in 6- well plates so that infections occurred at approximately 75% confluency. One hour before infection, the media was changed to Dulbecco’s modified Eagle’s medium (DMEM) (4.5 g/mL glucose). The bacterial inoculum was prepared by first growing EHEC strains statically in LB overnight at 37°C to OD600 of 0.6. Bacteria were resuspended in high-glucose DMEM to OD 0.5 and 45 uL of each inoculum was added to 5 wells of HT29 (MOI 10:1); DMEM alone was added to 5 wells as a mock infection (uninfected). The infections were carried out for a total of 6 hours, but the cells were washed once with DPBS and the media was replaced after 3 hours. After the 6- hour infection, each well was washed twice with DPBS to remove serum-containing media. RNA was extracted using the RNeasy Plus Mini Kit (Qiagen) and cDNA was generated from 2ug RNA using a High-Capacity cDNA Reverse Transcription kit (Thermo Fisher). Quantitative real-time PCR was performed using a Step One Plus Real-Time PCR machine using Taqman 2x master mix and Taqman probes for *GAPDH* (Hs02758991_g1), *IL23a* (Hs00372324_m1), *F3* (Hs00175225_m1) and *CXCL8* (Hs00174103_m1). Undiluted cDNA was used in the qPCR reactions. Expression levels were calculated using the delta-delta CT method normalized to *GAPDH*. Expression was normalized to the average expression of the 5 uninfected wells. Expression levels were compared using a two-tailed Student’s t-test. P-values were considered significant at less than 0.05 (*), 0.01 (**), 0.001 (***), and 0.0001 (****).

## Acknowledgments

We are grateful to members of the Waldor lab for comments on this project and the manuscript. We thank Dana-Farber/Harvard Cancer Center for the use of the Rodent Histopathology Core, which provided H&E staining of FFPE tissue sections and blinded histopathology scoring by Dr. Roderick T Bronson. Dana-Farber/Harvard Cancer Center is supported in part by an NCI Cancer Center Support Grant # NIH-5-P30-CA06516.

We thank the Harvard School of Public Health Biostatistics Student Consulting Center and the Microscopy Resources on the North Quad (MicRoN) core at Harvard Medical School (particularly Dr. Paula Llopis) for advice on appropriate statistical tests and quantitative image analysis.

MKW is funded by NIH grant R01-AI-042347 and the Howard Hughes Medical Institute.

## Supplementary Information

**Fig S1:**
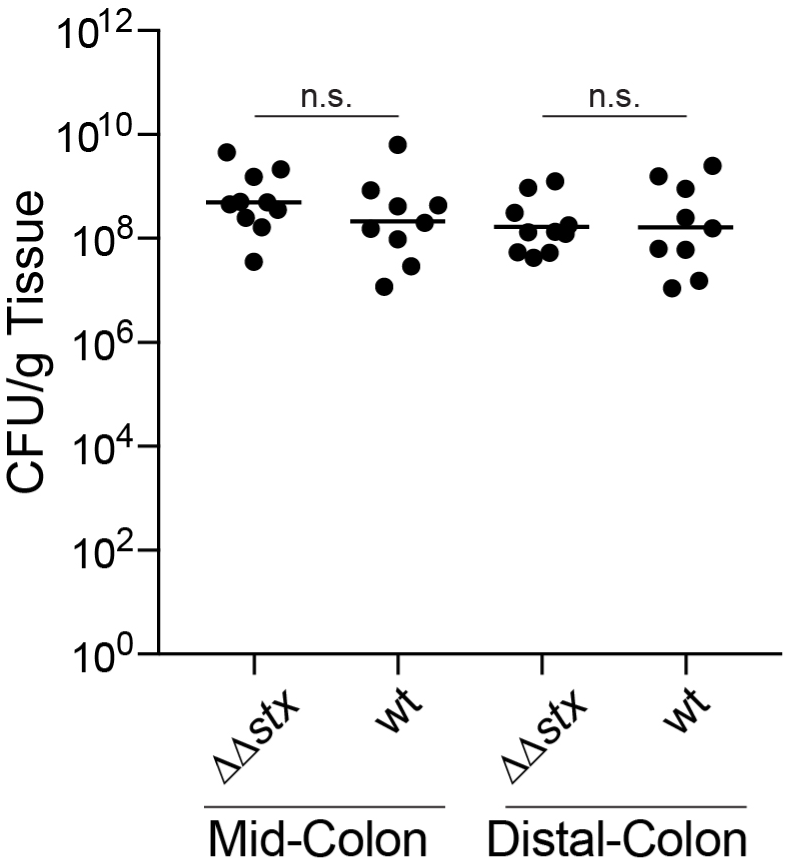
Intestinal colonization of WT and ΔΔ*stx* EHEC are similar. CFU recovered from mid or distal rabbit colon 36 hours post inoculation with either WT or ΔΔ*stx* EHEC. Lines indicate geometric mean. n.s. (not significant) by a two-tailed Mann-Whitney U statistical test.

**Fig S2:**
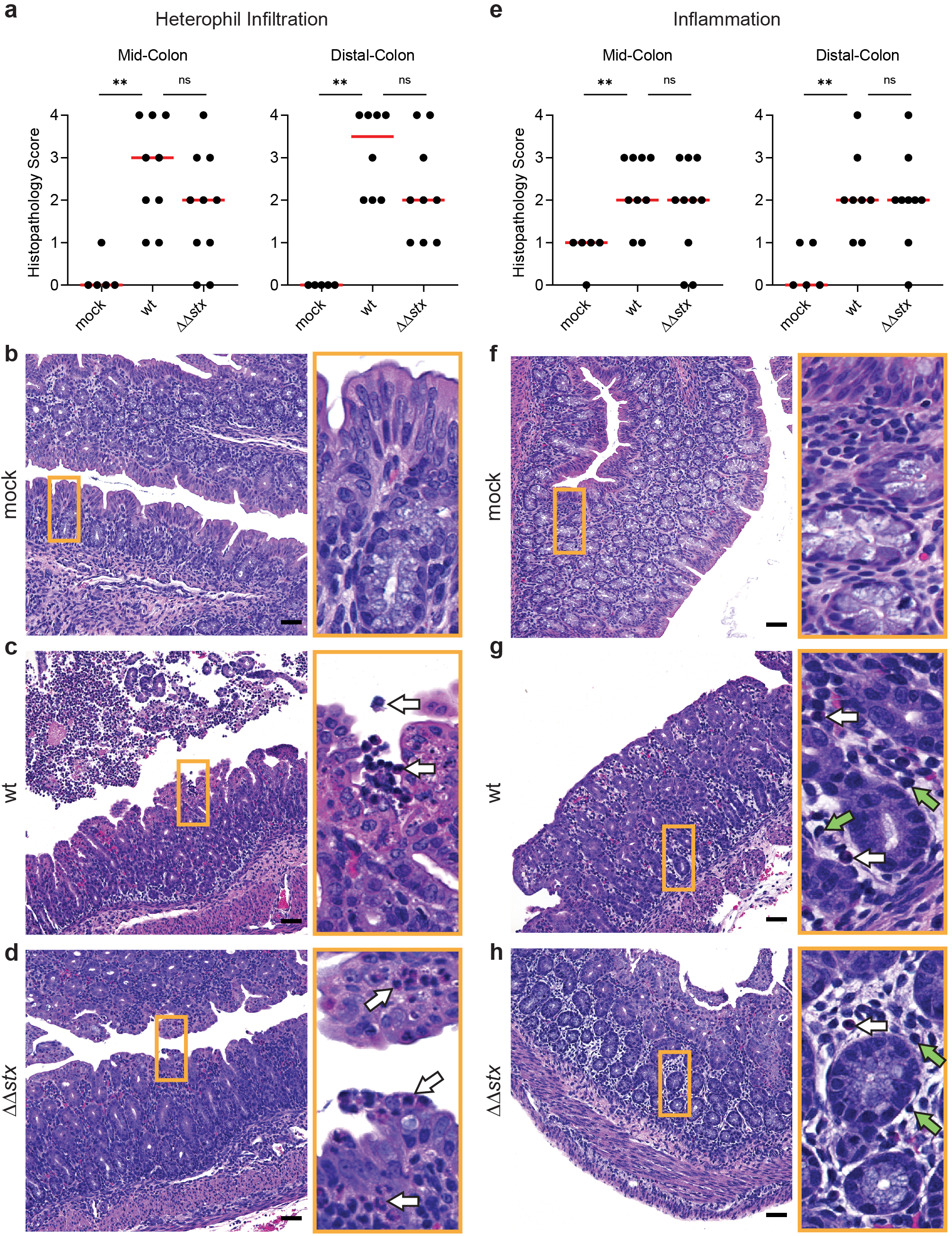
Immune cell infiltration is similar in animals infected with WT or ΔΔ*stx* EHEC. (a) Heterophil infiltration in colon sections from infant rabbits inoculated with PBS (mock), WT or ΔΔ*stx* EHEC 36 hours post inoculation. Scores for individual tissue sections are plotted along with the median (red line). Comparisons between groups was made using a two-tailed Mann-Whitney U test. P-values were considered significant at less than 0.05 (*) or 0.01 (**). n.s. indicates a non-significant difference. (b-d): Representative images from mock (score=0), WT (score=4), and ΔΔ*stx* (score=2) infected colons. Scale bars indicate 50 µm. Orange box denotes inset displayed to the right. White arrows indicate heterophils. (e): Severity of hemorrhage/edema. (f-h): Example images from mock (score=0), WT (score=3), and ΔΔ*stx* (score=3) EHEC-infected colons. Scale bars indicate 50 µm. Orange box denotes inset displayed to the right. White arrows indicate heterophils. Green arrows indicate lymphocytes.

**Fig S3:**
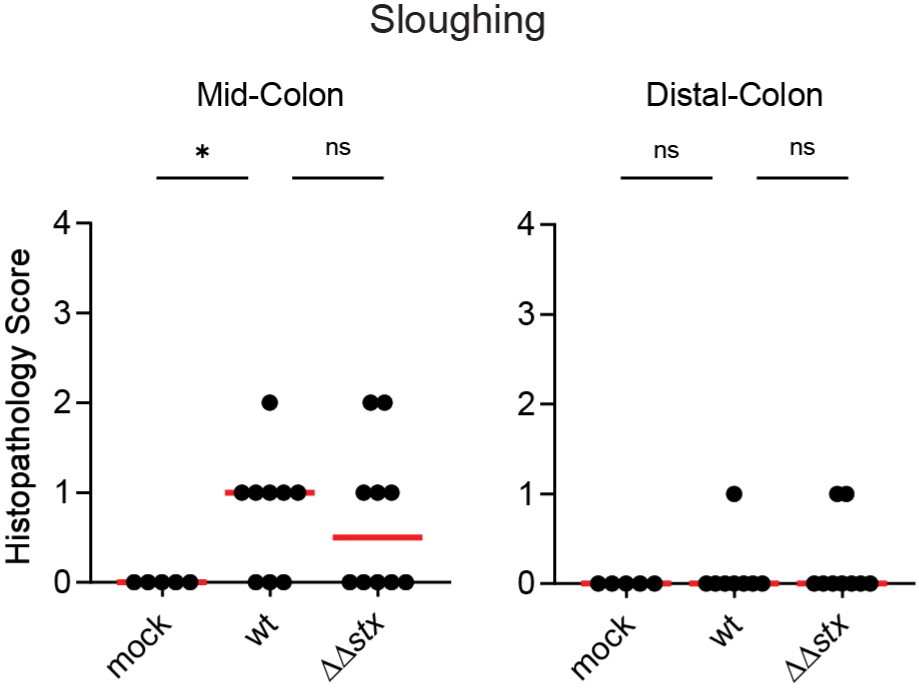
Epithelial sloughing is similar in animals infected with WT or ΔΔ*stx* EHEC. Sloughing in colon sections from infant rabbits inoculated with PBS (mock), WT or ΔΔ*stx* EHEC 36 hours post inoculation. Scores for individual tissue sections are plotted along with the median (red line). Comparisons between groups was made using a Mann-Whitney U test. The Bejmamini-Hochberg Procedure was used to control for the false discovery rate with multiple comparisons at 20%. P-values were considered significant at less than 0.05 (*). n.s. indicates a non-significant difference.

**Fig S4:**
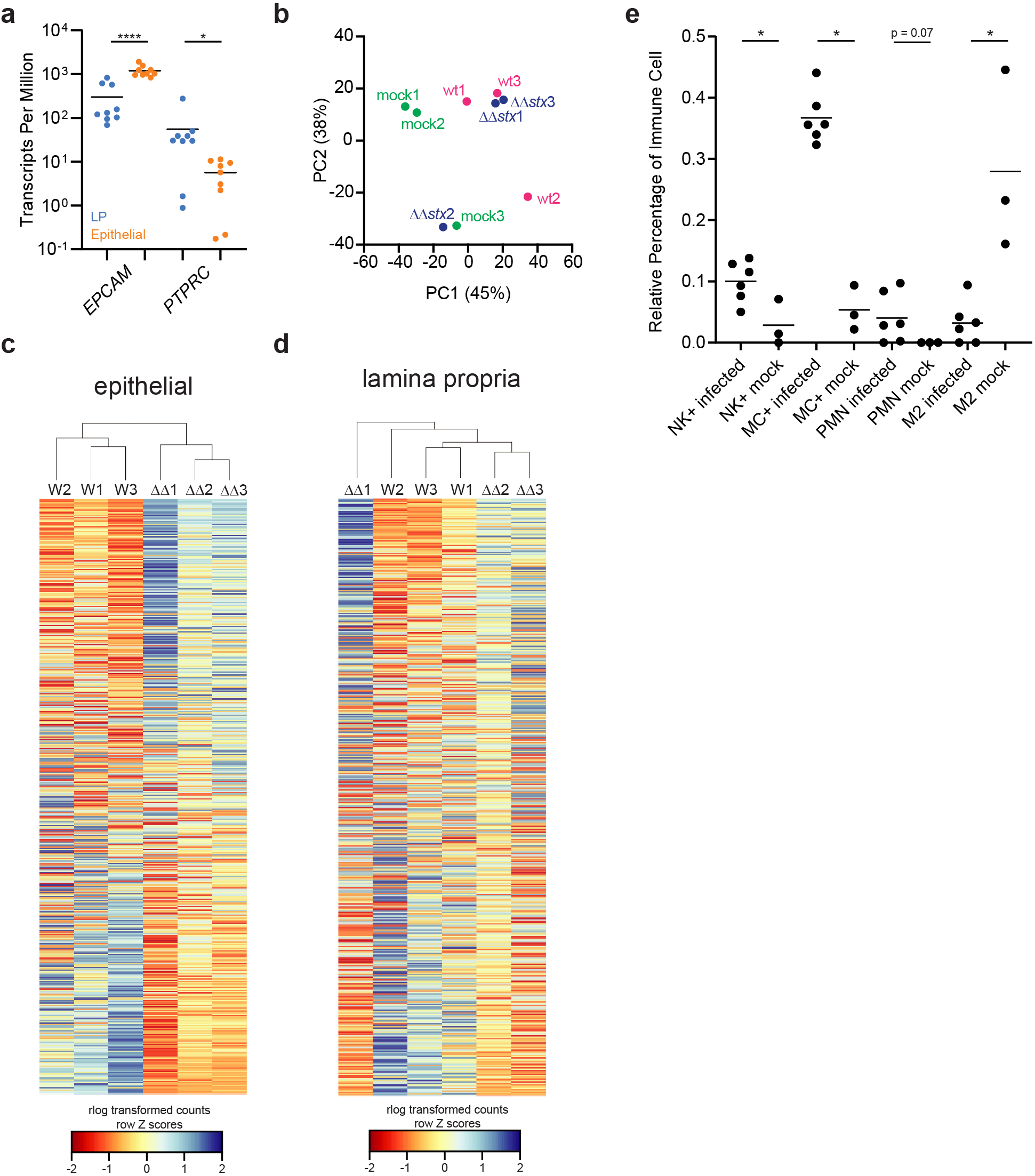
EHEC infection leads to transcriptomic changes in the colon. a) Relative gene expression in transcripts per million for epithelial cell marker *EPCAM* and immune cell marker CD45 gene *PTPRC.* Values for individual rabbits are plotted with mean. Distributions are compared with a Mann-Whitney U test, p< 0.05(*), 0.0001 (****). b) Principal Component Analysis (PCA) of rlog-transformed expression values of lamina propria cells. c) Heat map of rlog-transformed read counts from epithelial cells for 3 animal replicates (WT or ΔΔ*stx* EHEC infected) for all genes by rank. Hierarchical clustering performed using Euclidian sample distances. Rows are normalized by Z-score. d) Heat map of rlog-transformed read counts from lamina propria cells for 3 animal replicates (WT or ΔΔ*stx* EHEC infected) for all genes by rank. Hierarchical clustering performed using Euclidian sample distances. Rows are normalized by Z-score. e) Relative percentages of immune cell types in infected and uninfected tissue as predicted by CIBERSORTx. Distributions are compared with a Mann-Whitney U test, p< 0.05(*).

**Figure S5:**
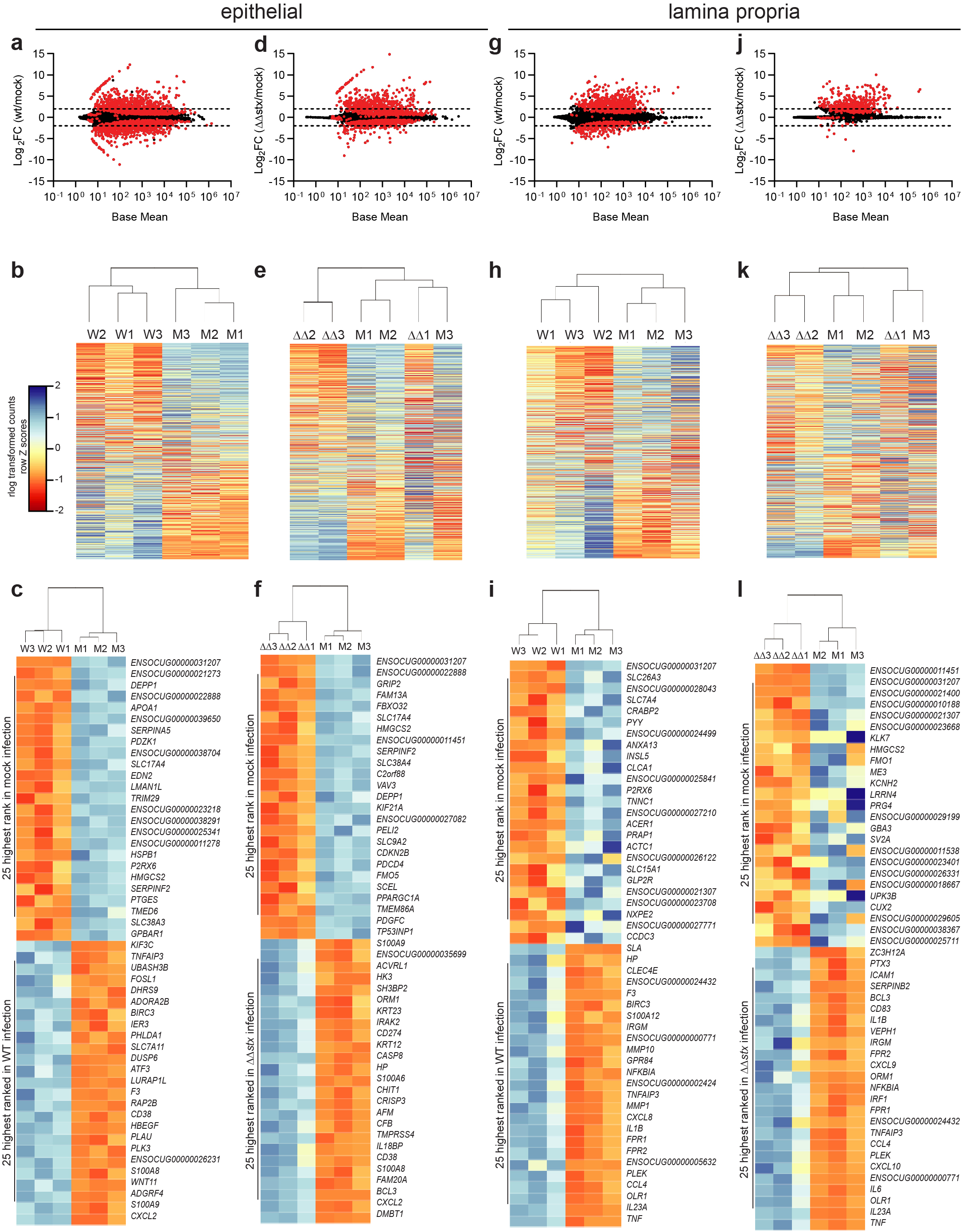
Comparison of transcriptional profiles from colonic samples of infant rabbits inoculated with PBS (mock), WT or ΔΔ*stx* EHEC. (a, d, g, j) Average expression level (base mean) and log2 fold change of transcript abundance in colonic epithelial cells (a, d) or lamina propria (g, j) from rabbits inoculated with WT EHEC (a, g) or ΔΔ*stx* EHEC (d, j) compared to PBS (mock). Genes with significantly different (adjusted p-value < 0.05) transcript abundance are highlighted in red. (b, e, h, k) Heat map of rlog-transformed read counts from epithelial cells for 3 animal replicates (WT or ΔΔ*stx* EHEC infected) for all genes by rank. Hierarchical clustering performed using Euclidian sample distances. Rows are normalized by Z-score. The 4 panels correspond to the comparisons shown in the panels immediately above (a,d,g,j); W, WT EHEC, M, mock, ΔΔ, ΔΔ*stx* EHEC. (c, f, i, l)) Heat map of rlog-transformed read counts from epithelial cells for 3 animal replicates (WT or ΔΔ*stx* EHEC infected) top 25 and bottom 25 genes by rank. The 4 panels correspond to the comparisons shown in the panels immediately above (b, e, h, k); W, WT EHEC, M, mock, ΔΔ, ΔΔ*stx* EHEC.

**Figure S6:**
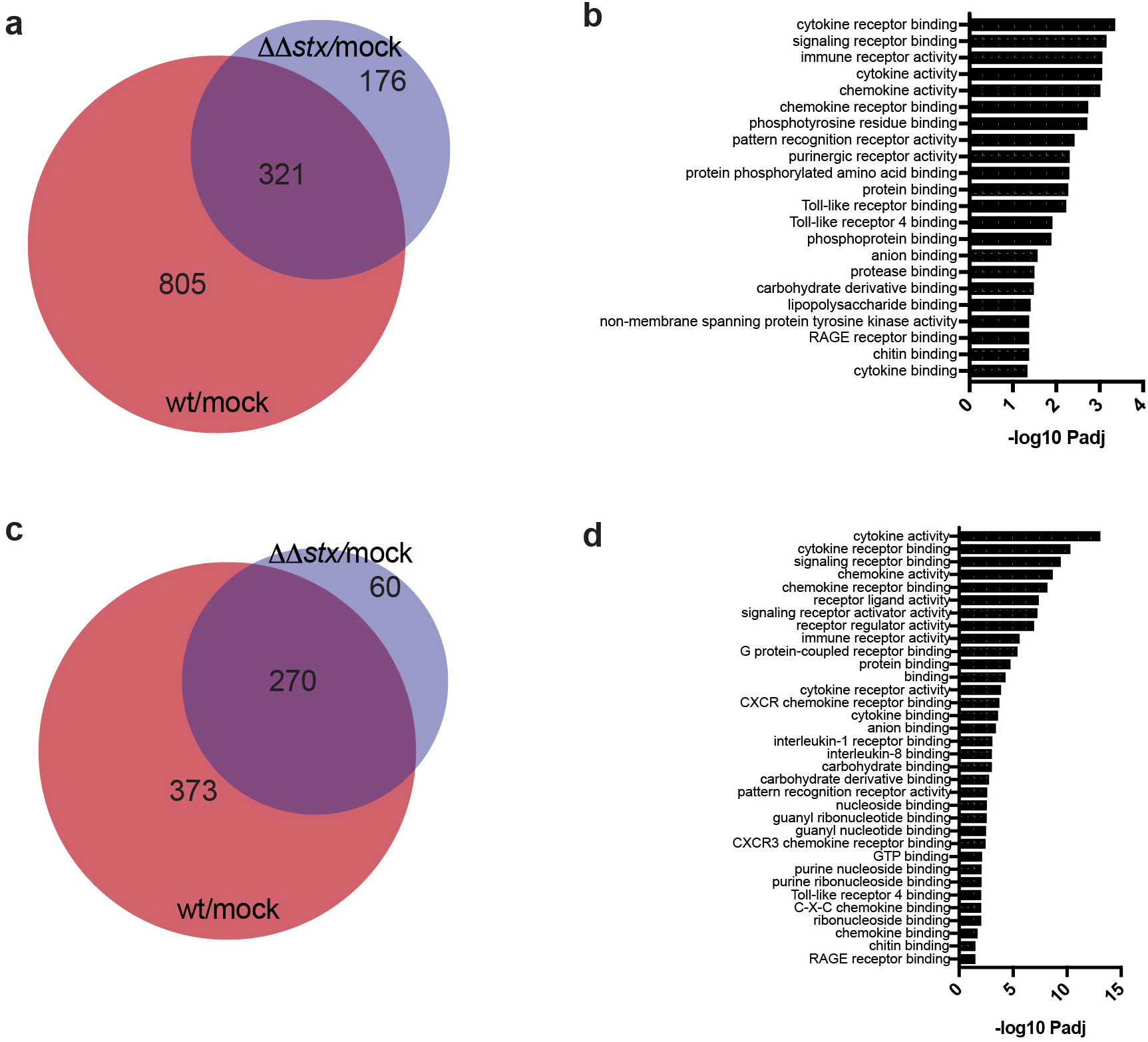
WT and ΔΔ*stx* EHEC colonic colonization both stimulate host transcriptional responses commonly associated with infection. a,c) Venn diagrams of differentially expressed genes in wt vs mock infection and ΔΔ*stx* vs. mock infection from colonic epithelial cells (a) or lamina propria cells (c). b,d) Transcriptional changes elicited by both strains map to many pathways associated with infection by GO Molecular Function analysis in colonic epithelial cells (b) or lamina propria cells (d).

**Figure S7.**
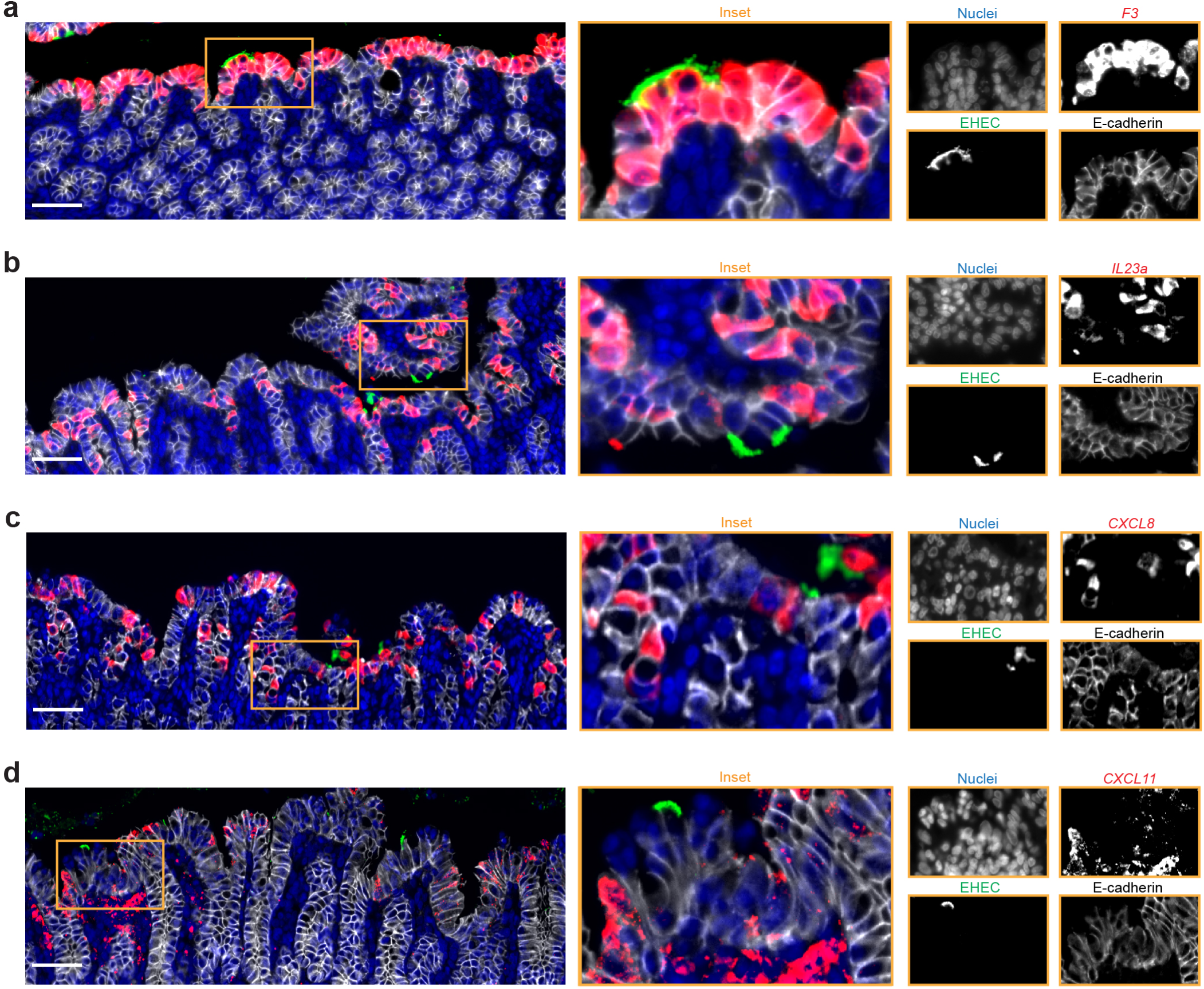
Bacteria do not colocalize with RNAscope signal. Immunofluorescence micrographs of colon sections from rabbits inoculated with WT EHEC (a-c) or ΔΔ*stx* EHEC (d) stained with an RNAscope probe (red) for rabbit *F3* (a), *IL23a* (b), *CXCL8* (c), or *CXCL11* (d), DAPI (blue), and an anti-E-cadherin antibody (white). Scale bar is 50 µM.

**Figure S8.**
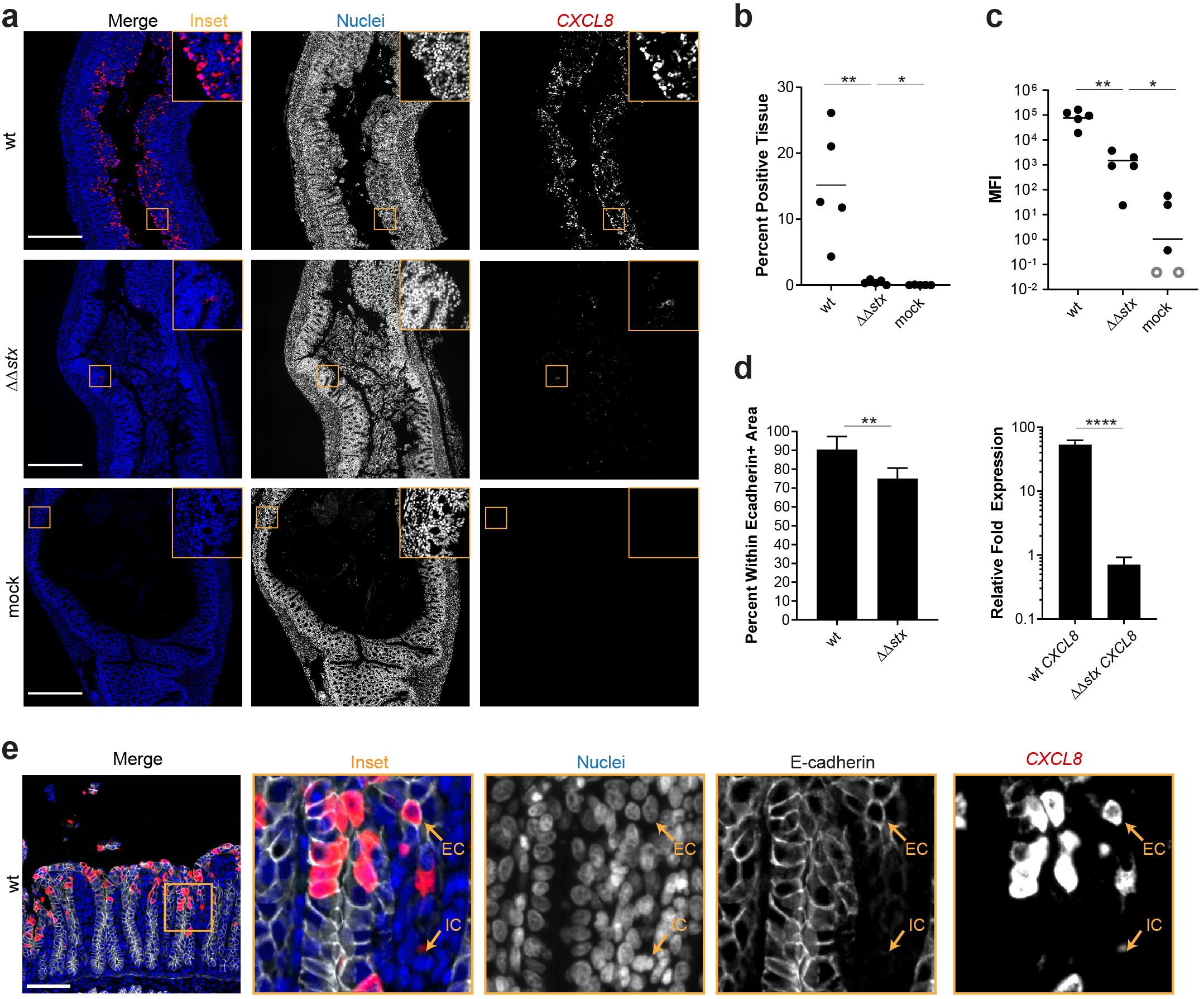
Expression of *CXCL8* in epithelial cells is much greater in animals infected with WT vs ΔΔ*stx* EHEC. a) Immunofluorescence micrographs of colon sections from rabbits inoculated with WT EHEC, ΔΔ*stx* EHEC, or PBS (mock) stained with an RNAscope probe for rabbit *CXCL8* mRNA (red) and DAPI (blue). Scale bar is 500 µM. b) Percentage of tissue section with *CXCL8* signal from individual rabbit colons. Distributions compared using the Mann-Whitney U test, p<0.01 (**), n.s. indicates not significant. c) Quantification of mean fluorescent intensity (MFI) of pixels above background from individual rabbit colons plotted with mean. Distributions compared using the Mann-Whitney U test, p<0.01 (**), n.s. indicates not significant. Open circles indicate limit of detection. d) Percent *CXCL8* signal within E-cadherin positive cells. Distributions compared using the Mann-Whitney U test, p<0.05 (*). e) Immunofluorescence micrographs of sections stained with an RNAscope probe to rabbit *CXCL11* mRNA (red), DAPI (blue), and anti-E-cadherin antibody (white). Scale bar is 500 µM. IC indicates immune cell, EC indicates epithelial cell. f) Normalized expression of *CXCL8* measured by RT-qPCR in HT29 colonic epithelial cells exposed to WT EHEC, ΔΔ*stx* EHEC, or PBS for 6 hours. Expression was normalized to the housekeeping gene *GAPDH* and then expressed relative to basal expression in uninfected cells. Statistical significance determined by a Students two-tailed t-test, p < 0.001 (***), n.s. indicates not significant.

**Dataset S1** Results of DESeq2 comparing WT vs mock infected rabbit colonic epithelial cells

**Dataset S2** Results of DESeq2 comparing ΔΔ*stx* vs mock infected rabbit colonic epithelial cells

**Dataset S3** Results of DESeq2 comparing wt vs mock infected rabbit lamina propria cells

**Dataset S4** Results of DESeq2 comparing ΔΔ*stx* vs mock infected rabbit lamina propria cells

**Dataset S5** Results of DESeq2 comparing wt vs ΔΔ*stx* infected rabbit colonic epithelial cells

**Dataset S6** Results of fGSEA comparing wt vs ΔΔstx infected rabbit colonic epithelial cells

**Dataset S7** Results of DESeq2 comparing wt vs ΔΔ*stx* infected rabbit lamina propria cells

**Dataset S8** Results of fGSEA comparing wt vs ΔΔstx infected rabbit lamina propria cells

